# *Salmonella* Enhances Osteogenic Differentiation in Adipose-Derived Mesenchymal Stem Cells

**DOI:** 10.1101/795617

**Authors:** Soraya H. Foutouhi, Nuradilla Mohamad-Fauzi, Dylan Bobby Storey, Azarene A. Foutouhi, Nguyet Kong, Amir Kol, Dori Borjesson, Prerak Desai, Jigna Shah, James D. Murray, Bart C. Weimer

## Abstract

The potential of mesenchymal stem cells (MSCs) for tissue repair and regeneration has garnered great attention. While MSC interaction with microbes at sites of tissue damage and inflammation is likely, especially in the gut, the consequences of bacterial association have yet to be elucidated. This study investigated the effect of *Salmonella enterica* ssp *enterica* serotype Typhimurium on MSC trilineage differentiation path and mechanism. Through examination of key markers of differentiation, immunomodulatory regulators, and inflammatory cytokines, we demonstrated that *Salmonella* altered osteogenic and chondrogenic differentiation pathways in human and goat adipose-derived MSCs. Gene expression profiles defined signaling pathway alterations in response to *Salmonella* association not observed in epithelial cells. We uncovered significant differential expression (*P* < 0.05) of genes associated with anti-apoptotic and pro-proliferative responses in MSCs during *Salmonella* challenge. These observations led us to conclude that bacteria, specifically *Salmonella,* induce pathways that influence functional differentiation trajectories in MSCs, thus implicating substantial microbial influence on MSC physiology and immune activity.

## Introduction

Application of mesenchymal stem cells (MSCs) to regenerative medicine is an area of intensive research [1, 2]. In addition to a capacity for self-renewal and differentiation into cartilage, bone, and adipose tissue [3], MSCs home to sites of inflammation where they exhibit immunomodulatory functions via secretion of paracrine factors [4, 5]. In cooperation with recruited immune cells, MSCs moderate inflammation via expression of anti-inflammatory cytokines [6–8], inhibit T-lymphocyte activation, and alter macrophages to express a regulatory anti-inflammatory phenotype to increase phagocytic activity [5, 9]. Bacterial infections promote inflammation and prompts MSCs to secrete anti-microbial peptides [10].

Microbial access to MSCs is likely at mucus membranes where tissue turnover is high and immune-responsive cells infiltrate to control pathogens [11]. Inflammation, and subsequent destruction of intestinal epithelial cells induces MSC recruitment to facilitate tissue recovery [12]. While an apoptotic epithelial cell response to infection is well characterized [13], little is known about the consequences of bacterial association on the behavior of stem cells. Treatment of MSC with lipopolysaccharide (LPS) and *Escherichia coli* increases osteogenesis and decreases adipogenesis; conversely, stimulation with *Staphylococcus aureus* decreases osteogenesis and adipogenesis [14]. Recent discovery of invasion in MSCs by numerous bacteria common in the oral cavity and gut resulted in augmentation of MSC inhibition of T-cell proliferation and provides evidence of direct alteration of MSC immune function [15]. Maintenance of viability [15] and change in differentiation path [14] confirms that bacterial association with MSCs not only varies in comparison to epithelial cells, but also altered function beyond acute infection.

Host detection of microbial presence is accomplished in part by Toll-like receptors (TLRs), which recognize bacterial components on their cell surface in epithelial cells are also expressed in MSCs [16, 17]. It is not clear how pathogens regulate these molecules in MSCs. Tomchuck et al. [18] reported the promotion of MSC migratory abilities, whereas a study by Pevsner-Fischer [19] found TLR activation shifts lineage commitment to proliferation.

*Salmonella* pathogenesis in epithelial cells is primarily mediated via the type three-secretion system [20, 21], which injects effector proteins that lead to apoptosis. These proteins target a variety of host cell regulators, including the NFκB pathway, leading to an inflammatory environment that increases microbial internalization [21, 22]. Early transcription factors implicated in MSC differentiation such as, peroxisome proliferator-activated receptor gamma (PPARG) [23] and secreted phosphoprotein 1 (SPP1) [24], are involved in inflammation, and may bridge cellular response to microbe-induced inflammation in MSCs. Previously unexplored differences between the infectious route in epithelial cells and the immunomodulatory changes in MSCs during host-pathogen interaction suggest substantial stem cell conditioning by *Salmonella*. These novel observations led our group to hypothesize that microbial modulation of the immune system in MSCs provides pathogens with an uncharted method of tissue infiltration, resulting in undescribed biological impact.

In this study, it was hypothesized that human and goat MSCs internalize *Salmonella* that resulted MSC altered trajectories towards a pro-osteogenic commitment, in parallel with the induction of an anti-inflammatory, immunosuppressive cellular phenotype. Examination of differentiation markers, immunomodulatory regulators and inflammatory cytokines demonstrated that *Salmonella* was internalized without inducing MSC apoptosis that altered osteogenic and chondrogenic differentiation. This phenomenon extended to alter molecular mechanisms of cell survival, proliferation and immune regulation. These observations found microbial-specific alterations in MSC differentiation and inflammatory status to influence stem cell fate and functionality.

## Materials and Methods

### Cell culture

Human adipose-derived mesenchymal stem cells (hASCs) were isolated by the laboratory of Dr. Dori Borjesson (University of California, Davis) and cultured in Minimum Essential Medium Alpha Modification (MEM-α, HyClone) with 20% fetal bovine serum (FBS, HyClone) and 1% penicillin-streptomycin (P/S, Gibco Life Technologies). Goat adipose-derived mesenchymal stem cells (gASCs) were isolated by the laboratory of Dr. Matthew Wheeler (University of Illinois, Urbana-Champaign), as described by Monaco et al. [25], and expanded as described by Mohamad-Fauzi et al. [26]. ASCs were cultured in 5% CO_2_/37°C, used at passage six. Colonic epithelial cells (Caco-2; ATCC HTB-37) were obtained from American Type Culture Collection (Manassas, VA) and grown as described by Shah et al. [27].

### Bacteria culture

*Salmonella enterica* ssp *enterica* serotype Typhimurium LT2 (ST), 14028S, serotype Enteritidis (BCW_4673), serotype Saint Paul (BCW_88) and serotype Newport (BCW_1378) were grown in Luria-Bertani (LB) broth (Teknova, Holister, CA) and incubated with shaking (200 rpm) at 37°C. Cultures were grown as described by Kol et al. [15].

### Quantification of microbe association

Association was determined using the gentamicin protection assay [28] and modified by Kol et al. [15], with the following modifications: ASCs were plated (4 x 10^4^) in a 96-well plate and incubated overnight; bacteria were suspended in serum-free medium (10^8^ CFU/ml) and added to the ASCs (MOI 100:1).

### Transmission Electron Microscopy

hASCs were plated on glass slides (Nalge Nuc International, Naperville, IL) and incubated for 2 h with ST [15]. Preparation and completion of transmission electron microscopy (TEM) was conducted as outlined by Kol et al. [15].

### Differentiation

Adipogenic and osteogenic differentiation was done using ASCs in 6-well plates at 2.5 x 10^5^ ASCs/well and incubated with ST for 1 h as described above. Chondrogenic differentiation was done in T-25 flasks at 3 x 10^5^ ASCs/flask. Following treatment with gentamicin, ASCs were washed with PBS and cultured for 48 hours in expansion medium to 70-80% confluence after which differentiation medium was added.

### Osteogenic differentiation assay

ASCs were cultured in osteogenic medium, fixed, rinsed and visualized under light and phase microscopy as described in Mohamad-Fauzi et al. [26]. hASCs were cultured for 14 days, whereas gASCs were cultured for 21 days. Control, non-induced cells were cultured in expansion medium.

### Chondrogenic differentiation assay

Chondrogenic differentiation was carried out as described by Zuk et al. [29]. Following ST incubation, 70-80% confluent cells were trypsinized and suspended in expansion medium for 14 days and then processed and visualized as described by Mohamad-Fauzi et al. [26].

### Adipogenic differentiation assay

Cells were cultured for 21 days in adipogenic induction medium, fixed, stained, and visualized as described by Mohamad-Fauzi et al. [26].

### RNA extraction and cDNA synthesis

ASCs were flash frozen prior to RNA extraction. For analysis of immunomodulatory factors, ASCs were plated in 6-well plates (3 x 10^5^ cells/well) and incubated with ST as described above. Treatment with LPS (Sigma) was added at 10 ng/ml. MSCs were washed with PBS, and immediately lysed with TRIzol Reagent (Life Technologies). Total RNA was extracted as described by Mohamad-Fauzi et al. [26]. Total RNA (1 µg) was used for first-strand cDNA synthesis using SuperScript II Reverse Transcriptase (Life Technologies) and oligo-dT primers according to the manufacturer.

### Quantitative RT-PCR

Primers (Supplementary Tables 1-2) were designed using Primer3 if not obtained from references. All primers spanned exon junctions or included introns. mRNA expression was quantified using Fast SYBR Green reagent (Life Technologies) on the Bio-Rad CFX96 platform (95°C for 20 seconds, 40 cycles of 95°C for 3 s and 60°C for 30 s), followed by melt curve analysis. Gene expression was normalized to GAPDH using 2^-ΔΔCT^ [30, 31]. Differences in differentiation gene expression were calculated as fold-changes relative to cells cultured in expansion medium (non-induced) and not treated with bacteria (non-treated). Inflammatory gene expression was calculated as fold-changes relative to non-treated control cells. Treatments were analyzed in pairwise comparisons using the Student’s t-test on the software JMP (SAS Institute) (*p ≤* 0.05). Data are presented as mean ± SEM with three biological and technical replicates.

### GeneChip expression analyses

Caco-2 infection samples with *Salmonella* LT2 were conducted using Affymetrix HGU133Plus2 GeneChip. Custom arrays containing all annotated coding and intergenic sequences of *Salmonella enterica* spp. *enterica* sv Typhimurium LT2 [32–34]. Data were normalized using MS-RMA [35] and analyzed using Significance Analysis of Microarrays (SAM) [33, 36].

### hASC RNA sequencing

Total RNA (1 µg) from hASCs was used to construct sequencing libraries with the Truseq Stranded Total RNA LT Kit (Illumina). Quality of RNA and constructed libraries was determined via 2100 Bioanalyzer. Libraries sequenced using an Illumina HiSeq2000 (BGI@UC Davis, Sacramento, CA) with single-end 50 bp. Reads were aligned using the UCSC hg19 human reference genome (ftp://igenome:G3nom3s4u@ussd-ftp.illumina.com/Homo_sapiens/UCSC/hg19/Homo_sapiens_UCSC_hg19.tar.gz) and annotated using “-a 10 --b2-very-sensitive -G”. Read counts and normalization was done using Cufflinks package (version 2.2.0) with flags “-u -G”. Tables from cuffnorm and cuffdiff imported into Ingenuity Pathway Analysis (IPA; Ingenuity Systems, version spring 2014). Sequence quality was examined using Phred [Supplementary Figure S1A-B].

### Ingenuity Pathway Analysis

IPA was used to determine biological pathways associated with gene expression profiles. Networks represent molecular interaction based on the IPA knowledge database. Estimation of probable pathway association was determined Fisher’s exact test, and predicted direction change was decided by the IPA regulation z-score algorithm (z-score ≥ 2 and ≤ 2 means a function is significantly increased or decreased, respectively) [37].

## Results

### Microbial association with adipose-derived mesenchymal stem cells

Human and goat ASCs were susceptible to *Salmonella* infection *in vitro* [Figure 1], which recapitulated previous observations in canine ASCs [15]. Total associated bacteria in both organisms were invasive; gASCs showed significantly higher invasion compared to human cells (*P* = 0.006) [Figure 1A]. Microbial association was not exclusive to ST; other *Salmonella* serotypes also invaded ASCs, as did other organisms as our group demonstrated previously [15], signifying a consistent trend in ASC vulnerability to common pathogens [Figure 1B]. Utilizing TEM, intracellular ST were observed 2 h post hASC-microbe co-incubation [Figure 1C-F]. Cells were not morphologically distressed nor apoptotic. In addition to invasion, ST adherence to hASC cell surface was observed; this intimate association is consistent with other non-pathogenic bacteria that Kol et al. [15] also observed.

**Figure 1:**
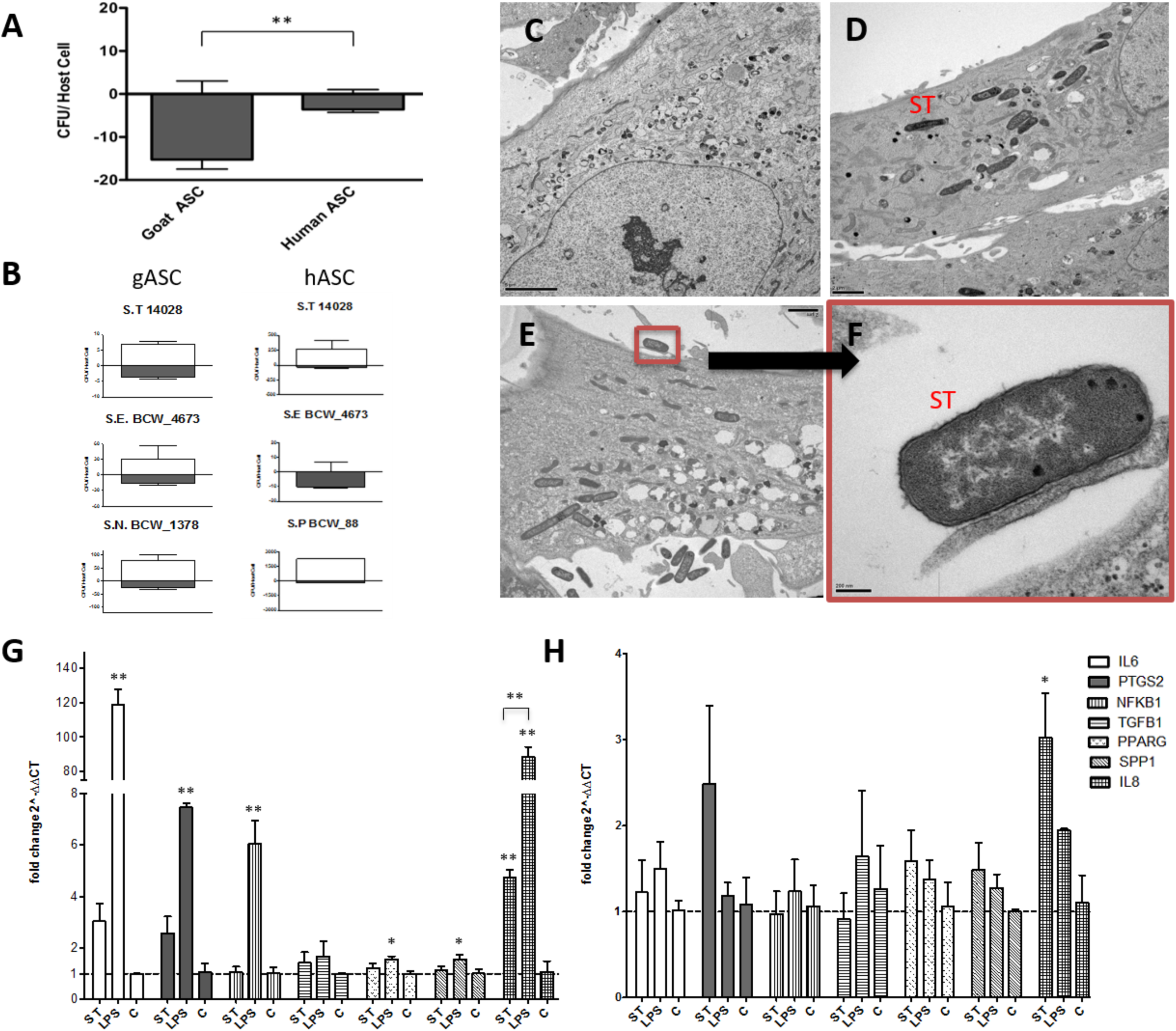
**Microbial Association with human and goat ASCs (A-F).** ASCs presented a uniform pattern of ***Salmonella enterica* ssp *enterica* serotype Typhimurium LT2 (ST)** infection, the total associated bacteria were invaded, gASC show significantly higher invasion compared to human cells (A). ASCs susceptibility to invasion was not exclusive to ST, association patterns were microbe specific; 35%, 12% *Salmonella enterica* ssp *enterica* serotype Typhimurium 14028, and 25%, 100% *Salmonella enterica* ssp *enterica* serotype Enteritidis (BCW_4673) were invaded in goat and human ASCs respectively (B). In gASCs, 35% *Salmonella enterica* ssp *enterica* serotype Newport (BCW_1378) and in hASCs, 7% of *Salmonella enterica* ssp *enterica* serotype Saint Paul (BCW_88) were invaded (B). Intracellular ST was observed by TEM two hours post MSC co-incubation (D-F), consistent with control non-treated hASCs (C), ST infected cells showed no signs of cellular toxicity (D-F). ST adherence to hASC was observed at various sites (E-F). **Expression of immunomodulatory factors in ASCs post-microbial association (G-H).** Quantitative PCR analysis of IL6, PTGS2, NFKB1, TGFB1, PPARG, SPP1, and IL8 expression in (G) goat and (H) human ASCs treated with ST or LPS. Data is presented as fold change (± SEM) in relative to expression levels in non-treated cells (“C”) (fold change ∼1, indicated by the dotted line). Statistical significance of *P <* 0.05 is denoted by an asterisk (*), and *P* < 0.01 denoted by two asterisks (**).

As ST is the most common cause of enteric diarrhea, we limited additional studies to this serotype. Following ST co-incubation, the expression of several immunomodulatory genes was investigated [Figure 1G-H]. Interleukin 8 and 6 (*IL8*, *IL6*), prostaglandin-endoperoxide synthase 2 (*PTGS2*), nuclear factor of kappa light polypeptide gene enhancer in B-cells 1 (*NFΚB1*), transforming growth factor beta 1 (*TGFB1*), *PPARG* and *SPP1* were selected. gASCs and hASCs treated with ST significantly increased *IL8* expression (*P ≤* 0.033, *P ≤* 0.005, respectively). An increase in *IL8* was also observed in LPS-treated gASCs (*P ≤* 0.0001). LPS-stimulated gASCs induced a significant increase in *IL6* (*P =* 0.0001), *PTGS2* (*P =* 0.0009), *NFKB1* (*P =* 0.0002), *PPARG* (*P* = 0.0204), *SPP1* (*P =* 0.037) and *IL8* (*P* ≤ 0.0001) gene expression.

A broader analysis of gene expression using RNAseq found 118 significantly differentially expressed genes in ST/hASCs interactions (*P* ≤ 0.05, FDR = 0.1). Canonical pathway analysis found hASCs treated with ST repressed cell death and survival genes associated with apoptosis [Figure 2]. TNF/FasL pathway analysis [Figure S2] established the muted apoptotic response in hASCs compared to Caco-2 cells following ST infection. Consistent with viability post ST association, induced genes in hASCs included repression of apoptosis, promotion of proliferation and multipotency [Figure 3.1]. Expression of heat shock protein B6 (*HSPB6*) and MAP-predicted activation of *v-akt* murine thymoma viral oncogene homolog 1 (*AKT1*) indicate promotion of hASC survival. *HSPB6* inhibits apoptosis of murine tumor cells, and plays a role in cellular protection against oxidative damage [38, 39]. Activated in response to a variety of cues, *AKT1* helps mediate cell survival and clonogenic potential [40–43].

**Figure 2:**
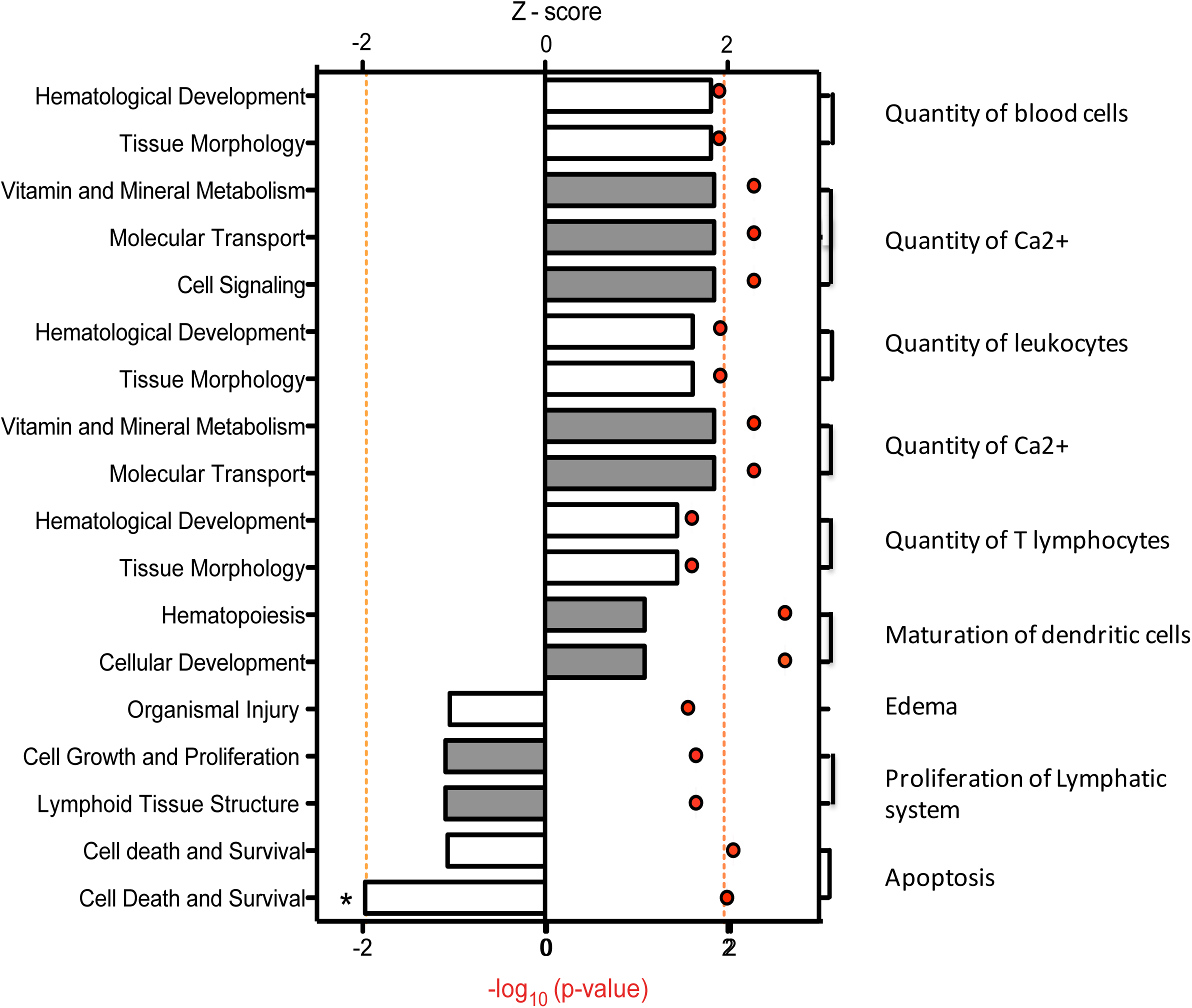
**Downstream trends analysis of differentially expressed genes in hASCs post microbial challenge.** The IPA regulation z-score algorithm was used to identify biological functions expected to increase or decrease based on the gene expression changes described in our dataset. Predictions base on p-value and z-score; positive z-score implies an increase in the predicted function, a negative z-score a decrease (z-score ≥ 2 or ≤ −2 represented by orange dotted lines). P-values ≤ 0.05 (red dots determined by Fischer’s exact test), illustrate a significant association between a given biological function and genes differentially expressed in our dataset (*P* ≤ 0.05).

**Figure 3.1:**
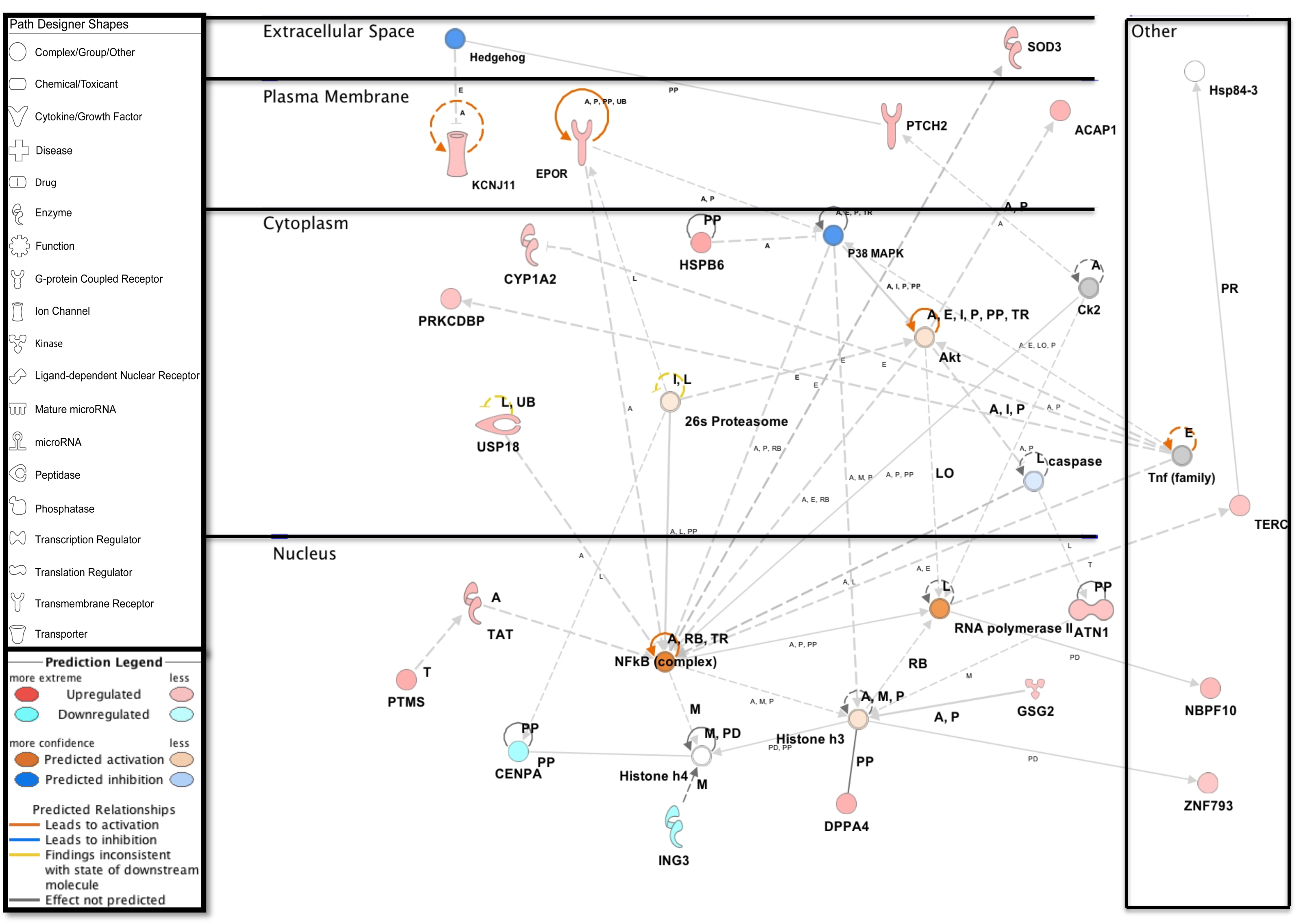
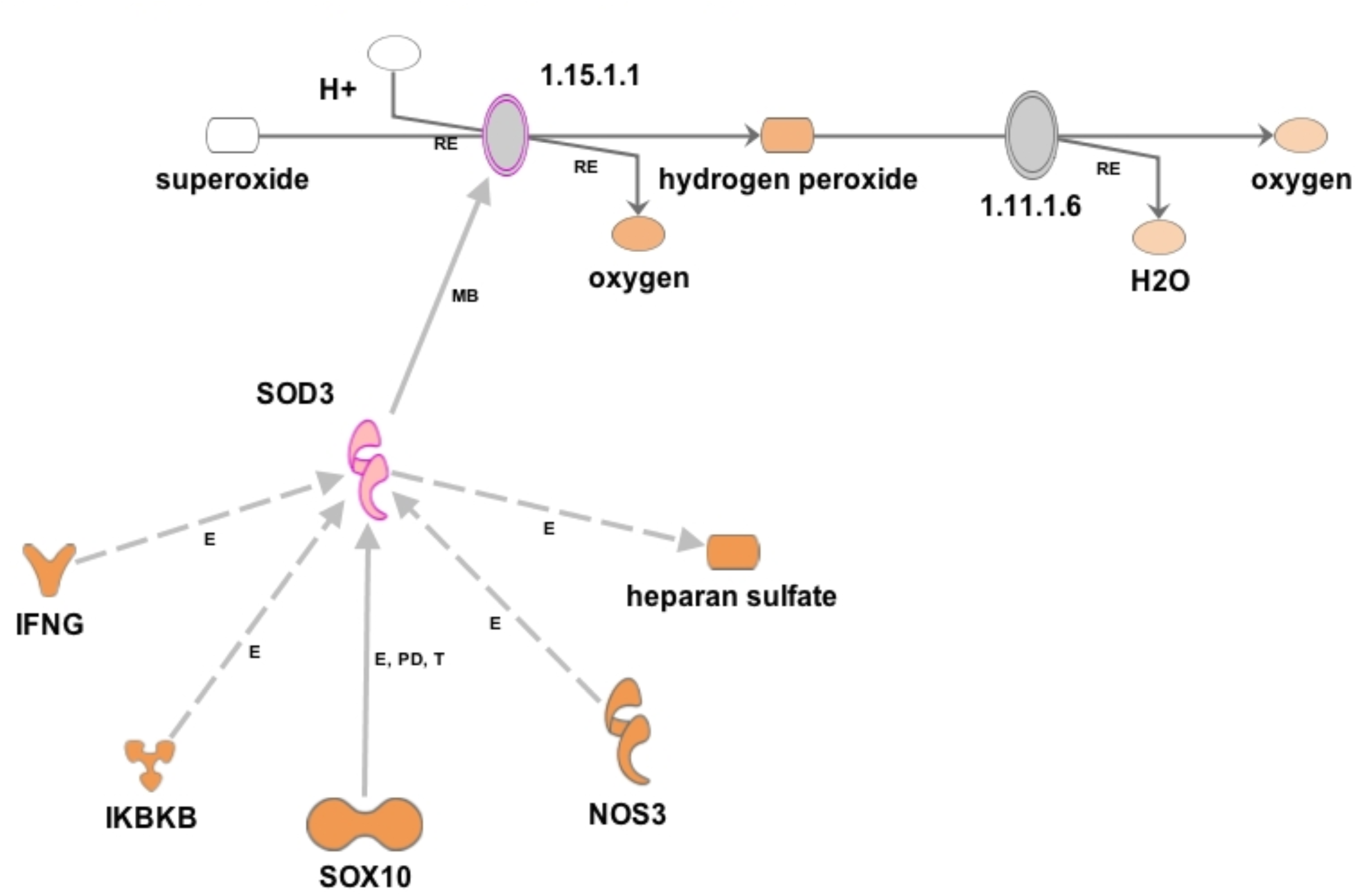
**Network displays interactions between genes regulating cell cycle, cellular assembly and organization that were differentially expressed in hASCs treated for sixty minutes with ST compared with untreated control.** Upregulated genes are colored in shades of red, down regulated in shades of green (*P* ≤ 0.05). IPA inserted Genes in white because they are connected to this network; dashed and solid lines denote indirect and direct relationships between molecules. The IPA molecule activity predictor assessed the activity of molecules strongly connected to this network; blue and orange colored molecules are predicted to have decreased and increased activity, respectively.

Upregulated in ST-treated hASCs [Figure 3.1], parathymosin (*PTMS*) is involved in molecular organization, differentiation and proliferation; nuclear translocation of *PTMS* is indicative of a pro-proliferative state [44, 45]. MAP predicted the repression of hedgehog signaling (Hh), whose inhibition is reported to decrease MSC proliferation with no effect on differentiation capacity [46]. We observed an increase in expression of patched 2 (*PTCH2*), which influences epidermal differentiation and Hh activity, suggesting promotion of ASC proliferation [47].

Superoxide dismutase 3 (*SOD3*) is pivotal for management of cellular redox [48–50] and was induced in this study [Figure 3.2], which aligns with previous observations in INFγ/LPS-activated microglial cells [48]. By promoting phagocytosis, EC-SOD facilitates bacteria clearance and elicits an anti-inflammatory response to LPS induced inflammation and pulmonary infection [51–53]. Interestingly, while Caco-2 cells strongly induced expression of TLR signaling which facilitates the LPS response following pathogen challenge, this observation was not seen in hASCs [Figure S3]. Using MAP, we focused on upstream regulators of *SOD3*, which may have been responsible for its activation [Figure 3.2]. Both *SOX10* and heparin sulfate (HS), play a role in the maintenance of multipotency and self-renewal [54, 55]. Interferon gamma (*INFG*) influences the immunomodulatory effects of MSC, as *INFG*-activated MSCs suppress T-cells and provide the necessary signal for MSC immunosuppression [56].

**Figure 3.2:**
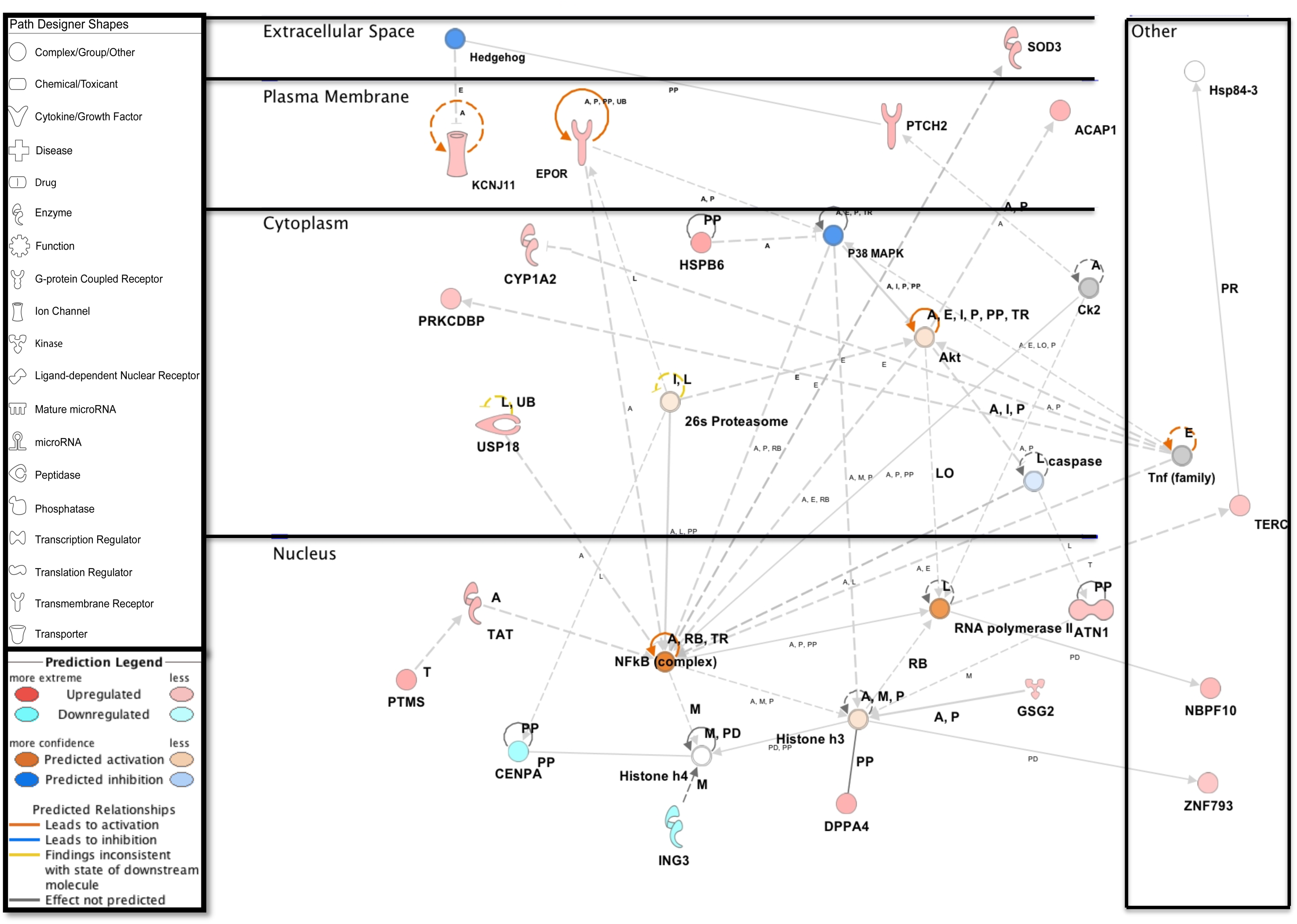
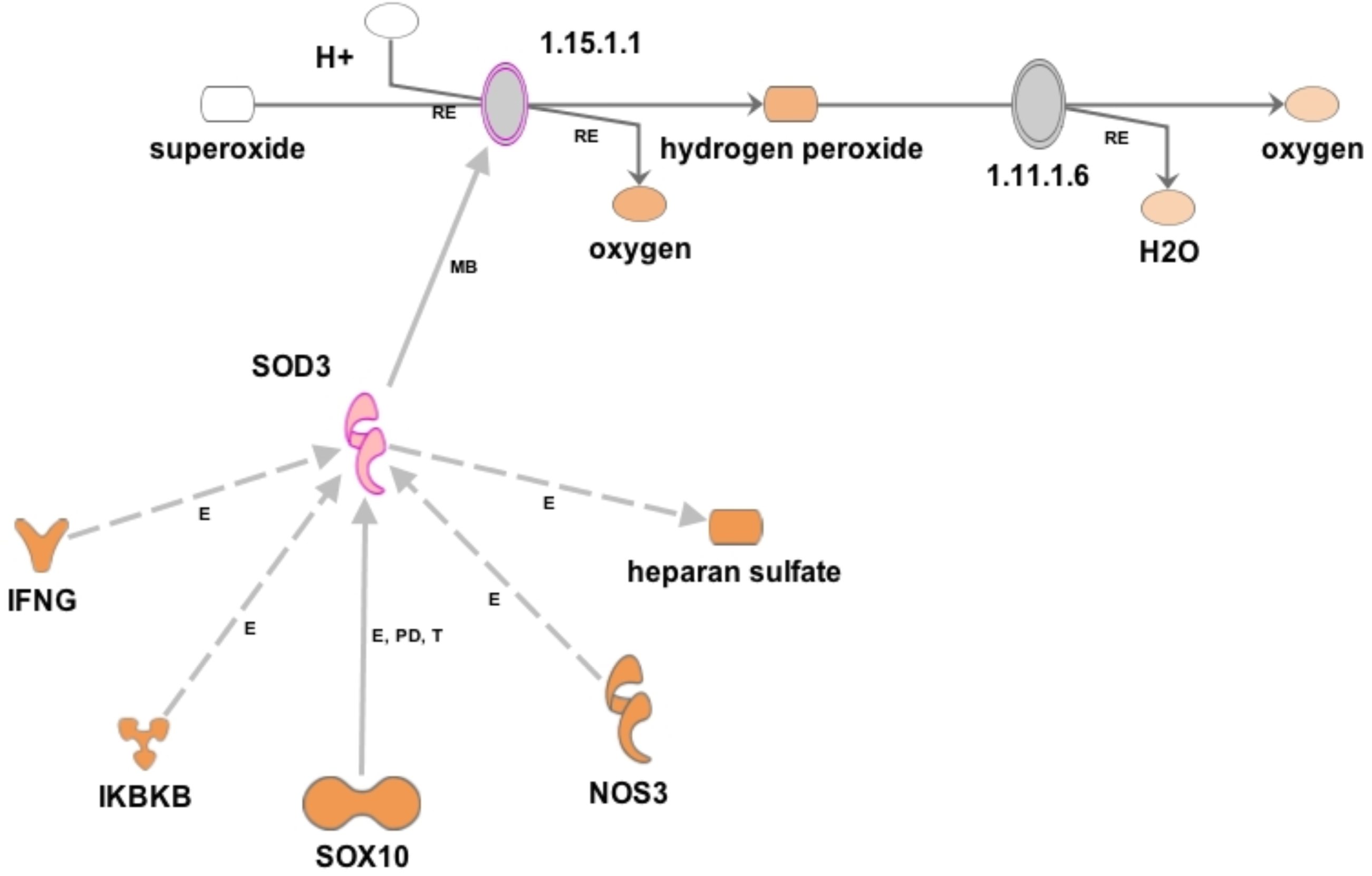
**Up regulation of superoxide dismutase 3 (SOD3) in hASCs following microbial challenge.** The IPA molecule activity predictor assessed the activity of molecules strongly connected to SOD3 (colored in shade or red); orange colored molecules are predicted to have an increased activity, based on the increased expression of SOD3 in our dataset. IPA inserted Genes colored in white because they are connected to this network; dashed and solid lines denote indirect and direct relationships between molecules.

### Analysis of trilineage differentiation post-microbial association Chondrogenic Differentiation

ST treatment did not abate ASCs ability to undergo chondrogenesis. In hASCs, differentiated cells migrated to form ridges that stained with Alcian Blue [Figure 4.2]. The morphological changes in gASCs were more advanced compared to human cells after ridge formation, cells aggregated, forming clumps that stained positive with Alcian Blue. Control, non-induced cells remained in monolayer and exhibited minimal background staining.

**Figure 4:**
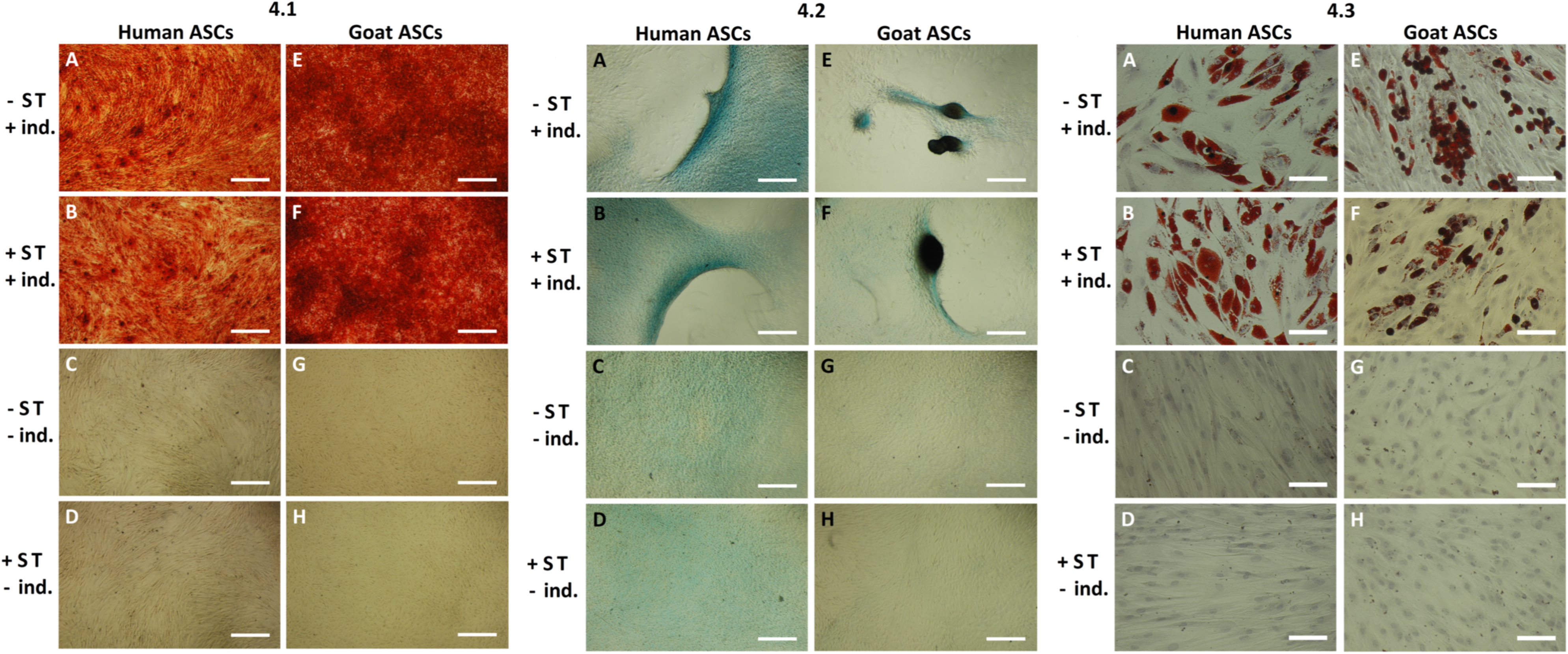
**Trilineage staining of human and goat ASCs.** Alizarin Red S staining of osteogenic differentiation in ASCs post-microbial association (4.1). hASCs (A-D) were cultured in osteogenic differentiation medium for 14 days, whereas gASCs (E-H) for 21 days and stained with Alizarin Red S. ASCs cultured in osteoinductive medium stained positive for calcium, (A-B and E-F, respectively), but did not stain when cultured in control medium (C-D and G-H, respectively), regardless of ST treatment. Representative images are shown in phase contrast at 40X magnification (scale bars represent 500 μm). Figure 4.2. Alcian Blue staining of chondrogenic differentiation in ASCs post-microbial association. Human (A-D) and goat (E-H) ASCs were cultured in chondrogenic differentiation medium for 14 days, and subsequently stained with Alcian Blue. Cellular condensation, as well ridge and micromass formations that stain positive was observed in human and goat ASCs induced for chondrogenesis (A-B and E-F, respectively), independent of ST treatment. Some background staining was observed in ST-treated and non-treated cells cultured in control medium, but cells remained in monolayer (C-D and G-H, respectively). Representative images are shown in phase contrast at 40X magnification (scale bars represent 500 μm). Figure 4.3 Oil Red O staining of adipogenic differentiation in ASCs post-microbial association. hASCs (A-D) and gASCs (E-H) were cultured in adipogenic induction medium for 21 days, and stained with Oil Red O. Accumulation of cytoplasmic lipid droplets were observed in ASCs induced for adipogenesis (A-B and E-F, respectively), independent of ST treatment. ST-treated and non-treated ASCs cultured in control medium did not yield lipid-positive cells (C-D and G-H, respectively). Representative images are shown in bright field at 200X magnification (scale bars represent 100 μm).

Expression of SRY (sex determining region Y)-box 9 (*SOX9*), which is essential for cartilage formation [57] and encodes for a transcription factor that promotes cartilage-specific extracellular matrix components [58, 59], was measured 14-days post chondrogenic induction. In hASCs, *SOX9* expression decreased (*P* = 0.04) in cells treated with ST [Figure 5.2]. Non-induced hASCs treated with ST showed no significant change in *SOX9* expression. Chondrogenic induction increased *SOX9* expression in hASCs compared to cells treated with control medium (*P* = 0.034). gASCs showed a significant decrease in *SOX9* expression in induced and non-induced ST-treated cells (*P* = 0.027, *P* = 0.039). There was a significant decrease in *SOX9* expression between gASCs treated with induction (*P* = 0.012).

**Figure 5:**
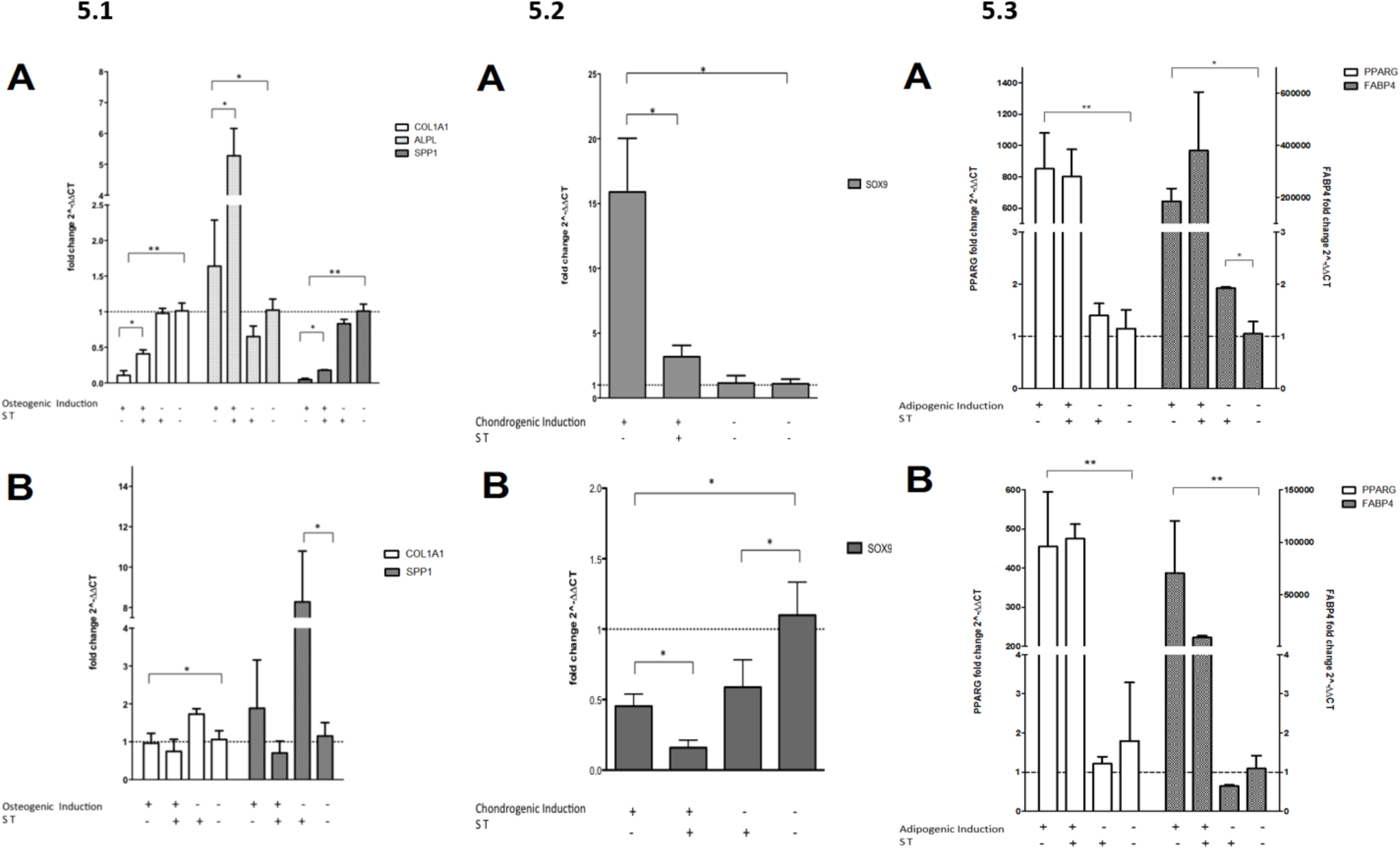
**ASC expression of trilineage differentiation markers.** Figure 5.1 Expression of osteogenic markers in ASCs post-microbial association. Quantitative PCR analysis of COL1A1, ALPL and SPP1 gene expression in A) human and B) goat ASCs induced with osteogenic differentiation medium and/or treated with S.T. Figure 5.2 Expression of chondrogenic markers post-microbial association. Quantitative PCR analysis of SOX9 expression in A) human and B) goat ASCs induced with chondrogenic differentiation medium and/or treated with S.T. Figure 5.3 Expression of adipogenic markers in ASCs post-microbial association. Quantitative PCR analysis of PPARG and FABP4 expression in A) human and B) goat ASCs induced with adipogenic induction medium and/or treated with ST. Data is presented as fold change (± SEM) relative to expression levels in non-treated, non-induced cells (fold change ∼1, indicated by the dotted line). Statistical significance of *P <* 0.05 is denoted by an asterisk (*), and *P* < 0.01 denoted by two asterisks (**).

### Adipogenic Differentiation

ASCs cultured in adipogenic medium accumulated lipid-filled vacuoles that stained with Oil Red O [Figure 4.3]. Visually, no differences were observed between non-induced ST-treated cells and the controls, which did not yield lipid-filled adipocytes nor stain with Oil Red O.

The lack of visual differentiation was further explored by examining expression of *PPARG* and fatty acid binding protein 4 (*FABP4*) 21 days post-induction, which should detect early events in adipogenesis [60, 61]. ST treatment of induced and non-induced hASCs did not change *PPARG* expression [Figure 5.3], a trend consistent with gASCs; however, a significant increase in *PPARG* expression in human and goat ASCs treated with differentiation vs. control medium (*P ≤* 0.0001, *P ≤* 0.0001) was observed, as expected.

*FABP4* is a fatty acid binding protein specific to mammalian adipose tissue [62, 63]. No significant difference was observed between induced ASCs treated with ST and non-treated control cells. Non-induced hASCs, but not gASCs, treated with ST displayed a significant increase in *FABP4* expression (*P =* 0.019). Both species induced *FABP4* in cells cultured in adipogenic induction medium [Figure 5.3] (*P =* 0.029, *P ≤* 0.0001).

### Osteogenic Differentiation

ASCs induced with osteogenic medium post ST treatment underwent osteogenesis. Mineralized calcium deposits accumulated within the monolayer and stained with Alizarin Red S [Figure 4.1]. By visual comparison, no difference was apparent between ST-treated and non-treated cells. ASCs cultured in control, expansion medium did not undergo calcium mineralization nor stain with Alizarin Red S, independent of ST treatment [Figure 4.1].

Collagen type I alpha 1 (*COL1A1*), alkaline phosphatase (*ALPL*) and *SPP1* expression were determined at the termination of differentiation. *COL1A1* encodes for the major component of the most abundant collagen found in bone matrix [64]. In hASCs induced for osteogenesis, *COL1A1* expression was significantly higher in ST-treated cells compared to controls (*P* = 0.025) [Figure 5.1A]. No difference in expression was detected between non-induced ST-treated and control hASCs. There was a significant decrease in *COL1A1* expression in hASCs treated with osteogenic induction vs. control medium (*P ≤* 0.0001). gASCs displayed no significant change of *COL1A1* expression in induced ST treated cells [Figure 5.1B]. There was a significant decrease in *COL1A1* expression between gASCs treated with osteogenic induction vs. control medium (*P* = 0.018).

*ALPL*, which provides phosphate ions for the production of bone mineral during matrix maturation [65–67], increased in induced hASCs (*P* = 0.02) as well as ST treated-induced hASCs. *ALPL* expression was 3.3-fold higher than in non-treated cells (*P* = 0.03) [Figure 5.1A]. In non-induced hASCs, no significant change in *ALPL* expression was detected following ST-treatment. *ALPL* expression in gASCs was not detected.

*SPP1* is a non-collagenous bone protein expressed during the mineralization phase late in osteogenesis [68]. In hASCs, *SPP1* was repressed in induced cells (*P ≤* 0.0001). *SPP1* expression was 4.3-fold higher in induced hASCs treated with ST compared to non-treated cells (*P* = 0.002). In non-induced hASCs, no difference in expression was observed between ST-treated and control cells [Figure 5.1A]. In induced gASCs, no significant change in SPP1 expression was observed in ST-treated cells. However, a 7.2-fold increase in SPP1 was observed in non-induced gASCs treated with ST compared to non-treated, non-induced controls (*P ≤* 0.05) [Figure 5.1B].

Pretreatment of hASCs with ST followed by 14 days of osteogenic induction resulted in the differential expression of 1060 genes (data not shown). RNAseq-IPA regulation z-score algorithm identified associated downstream biological functions. Expression data projected the repression of genes associated with cell-to-cell signaling, inflammation and response to infectious disease [Figure 6.1], highlighting the potential anti-inflammatory phenotype of hASCs subjected to microbial association (*P*-value ≤ 0.05, z-score ≥ 2).

**Figure 6.1:**
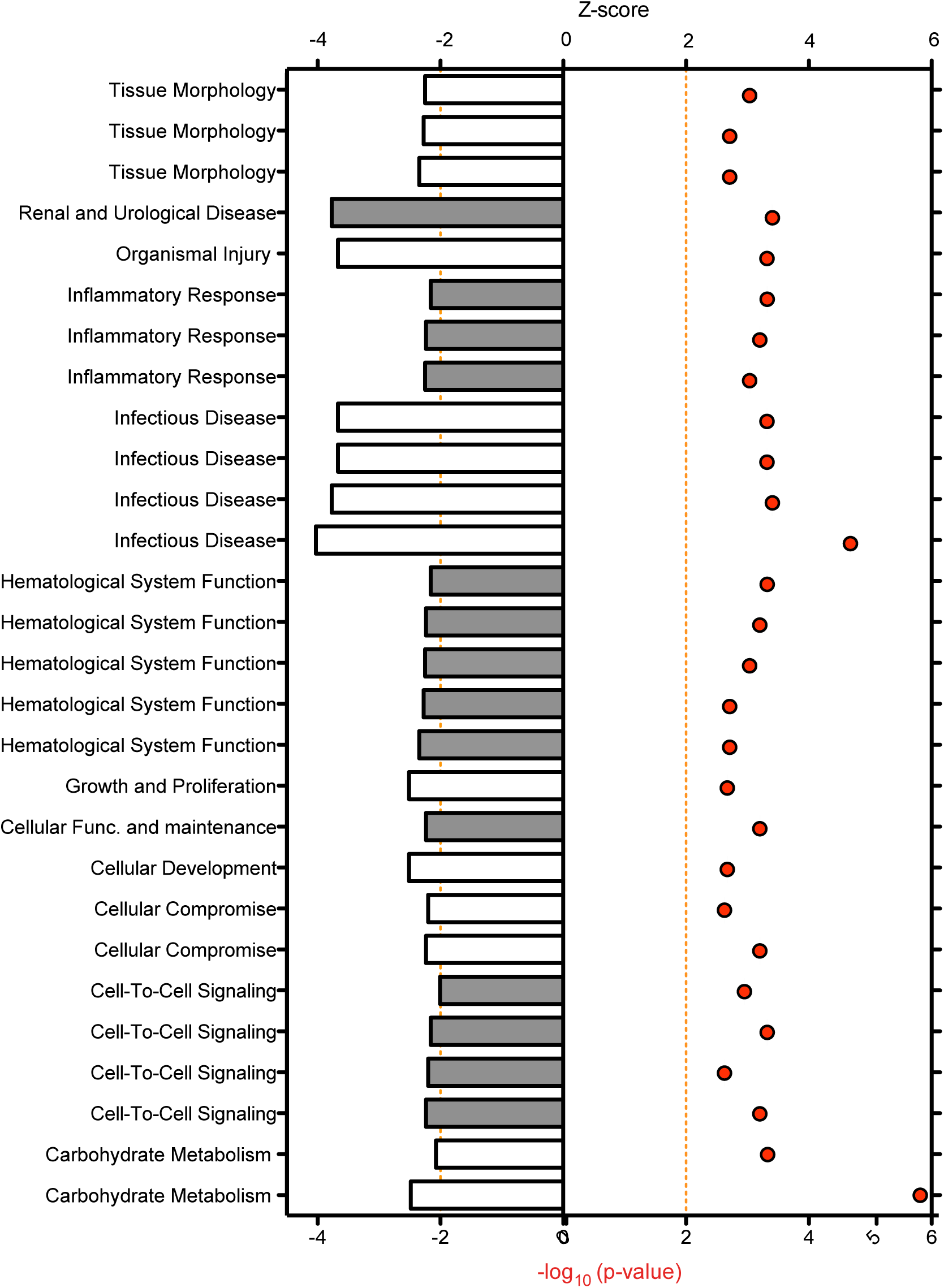
**Downstream trends analysis of differentially expressed genes in hASCs induced towards osteogenesis post microbial challenge.** The IPA regulation z-score algorithm was used to identify biological functions expected to increase or decrease based on the gene expression changes observed in our dataset. Predictions base on p-value and z-score; positive z-score implies an increase in the predicted function, a negative z-score a decrease (z-score ≥ 2 or ≤ −2 represented by orange dotted lines). P-values ≤ 0.05 (red dots determined by Fischer’s exact test), illustrate a significant association between a given biological function and genes differentially expressed in our dataset (*P* ≤ 0.05).

ST treatment followed by osteogenic differentiation also influenced genes involved in cellular communication, migration and lineage commitment. Differentially expressed genes included stanniocalcin 1 (*STC1*) and mesenchyme homeobox 2 (*MEOX2*) [Figure 6.2]. MSCs secrete *STC1* in response to apoptotic signals [69]. *STC1* is reported to play a role in the suppression of inflammation and may act in the regulation of calcium and phosphate homeostasis [69, 70]. *STC1* expression was downregulated in differentiated hASCs pre-treated with ST.

**Figure 6.2:**
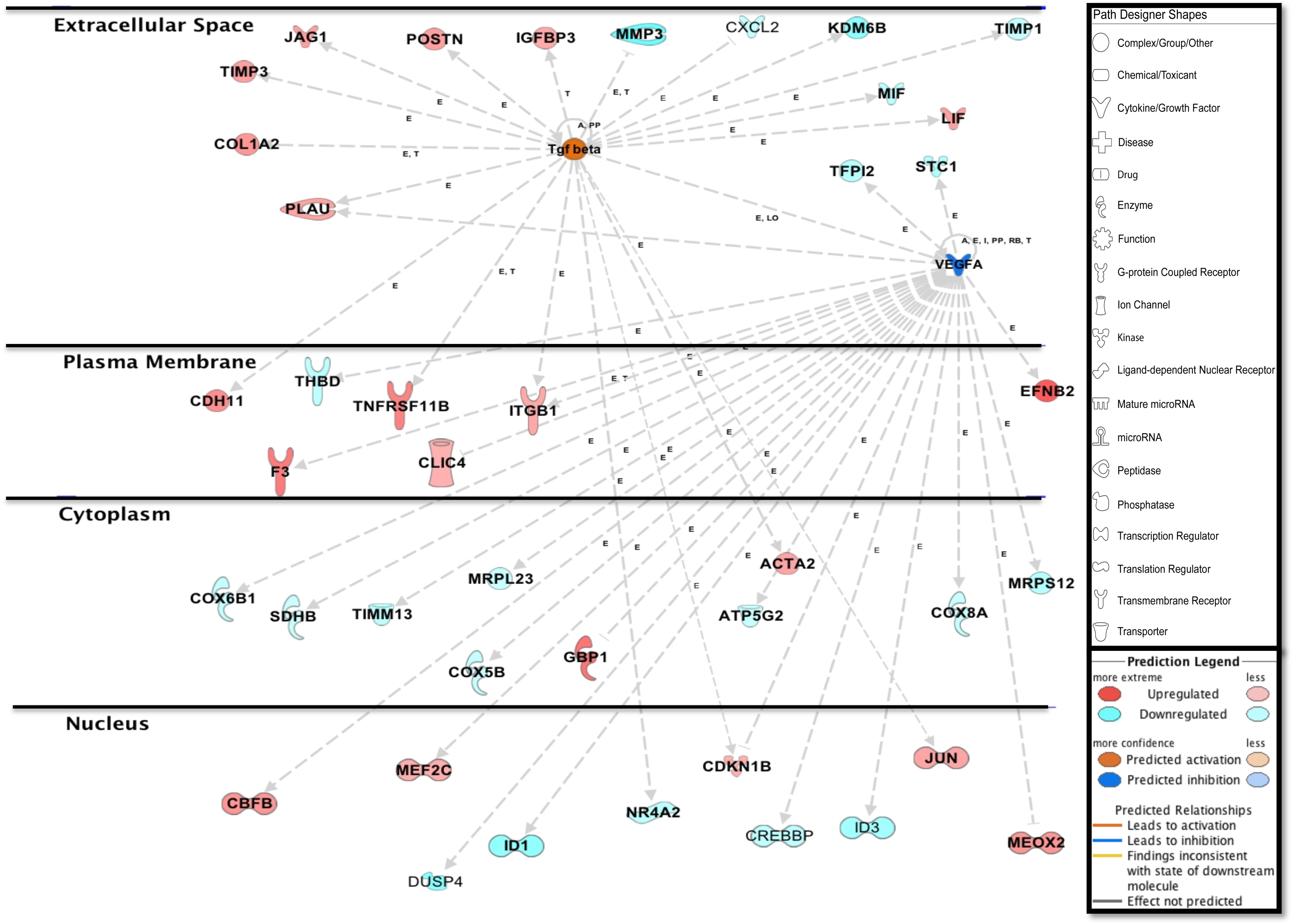
**Network displays interactions between genes involved in cell signaling, migration and differentiation that were differentially expressed in hASCs induced towards an osteogenic lineage following ST challenge.** Up-regulated genes are colored in shades of red, down-regulated in shades of green. Genes in white were inserted by IPA because they are connected to this network; dashed and solid lines denote indirect and direct relationships between molecules. The IPA molecule activity predictor assessed the activity of molecules strongly connected to this network; blue and orange colored molecules are predicted to have decreased and increased activity, respectively.

Upregulated in ST treated osteogenic differentiated hASCs [Figure 6.2], chloride intracellular channel 4 (*CLIC4*) is induced during cellular stress and influences cell cycle arrest and apoptosis [71]. Intracellular chloride regulates cation transport and may be involved in cellular signaling; *CLIC4* expression has been reportedly associated with Ca^2+^ induced differentiation of keratinocytes [71, 72]. Macrophage migration inhibitory factor (MIF) is elevated during tissue injury and inhibits MSC migration [73, 74]. Our data showed downregulation of MIF expression [Figure 6.2]. Furthermore, adhesion to fibronectin through α5β1-integrin plays a part in the induction of MSC migration [75]. MSC expression of A1-3, B1, and B3/4 integrins has been reported; blocking of B1 decreased migration in bone marrow-derived MSCs [76, 77]. We detected an increase in A3, A4 and B1 integrins [Figure S5].

Induction of periostin (*POSTN*) following ST challenge [Figure 6.2] was observed. A regulator of Wnt/β-catenin signaling cascade and mediator of bone anabolism, *POSTN* increases in response to stress and tissue damage [78, 79]. Wnt pathway activity is regulated by extracellular factors, including heparin sulfate proteoglycans, and acts to control MSC proliferation and differentiation [80]. hASCs treated with ST prior to induction displayed downregulation of Wnt activator Secreted frizzled-related protein 1 (*SFRP1*) and induction of transcription factors *JUN* and *AXIN2* [Figure S4A]. In addition, we observed a marked increase in the expression of *TNFRSF11B* [Figure S4B]. Also known as osteoprotegerin, *TNFRSF11B* is an anti-osteoclastogenic regulator, which is reported to augment osteogenesis [81].

Ephrin-B2 (*EFNB2*) is involved in osteogenic commitment and is required for differentiation of osteoclasts and osteoblasts *in vivo* [82]. Upregulated in our data set [Figure 6.2], the *EFNB2* ligand and *EPHB4* receptor are reportedly expressed on the surface of MSCs [82]. Increased expression of *COL1A2* was also detected [Figure 6.2]. *COL1A2* promotes cellular proliferation and osteogenesis, a response in part regulated by ERK/AKT1 pathway activation [83]. Activation of ERK mitogen activated protein kinase family (*MAPK*) drives the progression of osteoblasts via phosphorylation of transcription factors [77]. Overlay of hASC gene expression data illustrated upregulation of integrins involved in *MAPK1* activation as well as intracellular signal transducer phosphatidylinositol-4,5-bisphosphate 3-kinase (*PI3K*) [Figure S5].

Consistent with MSC ability to influence the immune response, we detected expression of genes involved in MSC signaling and immunomodulation [Figure 6.2]. Cadherin 11 (*CDH11*) expression was upregulated in our dataset. TGFB treatment increases expression of *CDH11* and subsequent calcium-dependent cell-to-cell interactions in MSCs [84]. Engagement of *CDH11* on fibroblast-like synoviocytes (FLS) has been reported to produce inflammatory mediators IL6 and IL8 [84], though our transcriptome analysis did not indicate significant upregulation of either of these cytokines. Kol et al. demonstrated that ST augments cMSCs ability to inhibit T-cell proliferation [15]. In hASCs, we observed the differential expression of genes involved in the suppression of immune cells; IPA regulator analysis highlighted the potential inhibitory effect of ST-treated hASCs on phagocyte and granulocyte proliferation [Figure S6].

## Discussion

Microbial presence on host tissue presents a wide range of beneficial to pathogenic effects on cellular function. Intestinal bacteria have been shown to contribute to cellular proliferation and development; observation of host and microbe interactions during inflammatory and disease states have been imperative in refining our understating of healing and self-renewal signaling mechanisms [85]. Studies on pathogen interference on MSC functionality demonstrate the promising potential of stem cells as a mode of intercession for infection and inflammation. Yuan et al [86] illustrated the ability of bone marrow-derived MSCs to increase clearance of methicillin-resistant *Staphylococcus aureus* (MRSA) in a rat model; work by Maiti and colleagues show that MSC stimulation with MRSA resulted not only in changes to cell proliferation but also the induction of inflammatory markers [87]. Here, ASCs were susceptible to microbial infection *in vitro* with many types of *Salmonella*, suggesting this is a common characteristic, especially when taken with the observations of Kol et al. [15]. Our specific investigation of ST resulted in changes in the expression of prototypical gene markers of differentiation and inflammation, but not apoptosis.

Fiedler et al. [14] found continuous treatment with heat-inactivated *E. coli* slightly increased ALPL activity in hASCs; heat-inactivation may have diminished the effect of bacterial exposure. The migration of MSCs to sites of inflammation, where bacterial interaction may be transient, could potentiate a narrow window of opportunity for microbial association. The use of viable bacteria and shorter exposure time in this study may better mimic the physiological conditions in which MSCs interact with microbes. We observed internalization of ST within 60 minutes of co-incubation; additional investigation is required to evaluate ST presence throughout extended culture periods. ST pre-treatment did not abate the ability of ASCs to differentiate, but did affect the expression of genes involved in osteogenesis and chondrogenesis.

ST treatment had a significant effect on osteogenic differentiation. In congruence with Fiedler et al. [14], we observed an increase in *ALPL* expression in ST-treated hASCs induced for osteogenesis. Increase in *ALPL* expression is consistent with the finding at 10 days post-osteogenic induction in LPS-treated hASCs [16]. *ALPL* was not detected in gASCs, as cells were harvested 21 days post-induction when the mineralization phase was likely occurring [88] and *ALPL* expression may have decreased [67, 88].

Consistent with the lack of *ALPL*, upregulation of *SPP1* (a late marker of osteogenesis) [88, 89], was observed in ST-treated gASCs. In addition to osteogenic commitment, the 6-glycosylated phosphoprotein SPP1 has a significant role in cellular stress and immunity. As an inflammatory mediator, SPP1 is reported as anti-inflammatory in acute colitis, while having an opposite effect in chronic disease status [90, 91]. Plasma levels of SPP1 are elevated in Crohn’s disease and SPP1-deficient mice have an impaired ability to clear *Listeria monocytogenes* [90, 92]. The upregulation of *SPP1* observed implies a diverse series of biological effect on MSC physiology. It is probable that the involvement of SPP1 in response to microbe-induced inflammation results in the influence of ASCs towards osteogenesis.

Differential marker expression following ST treatment was heavily observed in non-induced ASCs. Upregulation of *SPP1* was observed in ST-treated, non-induced gASCs, but not in induced cells. It is possible that the osteoinductive effect of ST treatment in induced ASCs was masked, as the medium contained additives that strongly induce osteogenesis. Pevsner-Fischer et al. [19] observed a differential effect of LPS treatment on non-induced MSCs compared to induced. Thus, the upregulation of osteogenic markers in non-induced cells highlight the direct influence of microbial treatment on ASC lineage commitment.

We observed a concomitant decrease in chondrogenic differentiation in response to ST treatment, as shown by a decrease in *SOX9* expression in ASCs. To our knowledge, this is the first report on the direct effect of bacterial association on MSC chondrogenesis. SOX9 is required for commitment to chondrogenic lineage, thus murine *SOX9*-null cells do not express markers of chondrogenesis [93]. Osteogenesis and chondrogenesis are tightly coupled processes [94], both regulated by proteins in the TGFβ superfamily [95]. An inverse relationship between osteogenic and chondrogenic differentiation has been demonstrated; microRNAs targeting genes important for osteogenesis were upregulated during chondrogenesis, and vice versa [96]. This further supports the premise that ASCs pre-exposed to ST prior to induction favor characteristics of osteogenic lineage commitment.

Exposure to whole bacteria and microbial components are sufficient to influence ASC signaling. MSC response to microbial components is mediated by Toll-like receptors (TLRs); hASCs express *TLR1-6* and *TLR9* [16]. TLR4 agonist LPS is a key component of the *Salmonella* cell wall [97]. LPS influences osteogenesis in hASCs and bone marrow-derived MSCs (BM-MSCs) by increasing mineralization, ALP activity, and expression of osteogenic markers [14, 16, 98, 99]. It is possible that LPS-induced changes in differentiation are mediated by TLRs and thus, dependent on NFKB1 activation [99, 100]. In this study, we did not observe upregulation of *NFKB1* expression following ST challenge or significant induction of genes involved in TLR signaling pathway. As a rapid responder, NFKB1 proteins are available and inactive; activity depends on phosphorylation-dependent degradation of NFKB1 inhibitors, thus the lack of change in mRNA expression is not unexpected [101].

While the conditions of this study did not lead to an observed induction of TLR gene expression, previous reports highlight the role of TLR activation in MSC physiology. TLR2 activation inhibited spontaneous adipogenic differentiation and increased osteogenesis in non-induced mouse BM-MSCs, but inhibits trilineage differentiation in induced cultures [19]. The upregulation of osteogenic markers in non-induced ST treated ASCs in this study supports this observation. Furthermore, TLR-activated MSCs recruit immune cells; TLR-activated macrophages secreted oncostatin M, a cytokine that induces osteogenic and inhibits adipogenesis in BM-MSCs [102]. MSCs deficient in myeloid differentiation primary response 88 (*MYD88*), which is crucial for TLR signaling [103], lack both osteogenic and chondrogenic potential [19], providing further evidence for linking microbe-induced TLR signaling and osteochondrogenic pathway induction.

Initiation of epithelial inflammation and rapid induction of pro-inflammatory cytokines via calcium-mediated activation of NFKB1 in response to *Salmonella* is documented [104, 105]. A dose-dependent increase in IL8 secretion was reported in human BM-MSCs following LPS treatment [106]. A hallmark of IL8 is the capacity for a variety of cells, including MSCs, to rapidly express and secrete IL8 [107]. IL8 can inhibit osteoclasts bone resorption activity, and stimulates osteoclast motility [108, 109]. Upregulation of *IL8* was detected after 14 days of osteogenic media treatment [108]. In additional support of the coordination between inflammatory and differentiation pathways, co-incubation with ST significantly increased *IL8* expression in hASCs and gASCs.

Coordination between inflammation and lineage commitment likely involves various small molecules. A pleiotrophic cytokine, IL6 is involved in innate tissue response to injury and maintenance of undifferentiated MSCs status, and *IL6* expression decreases during osteogenic differentiation [110]. While mature osteoblasts displayed enhanced osteogenic differentiation, primitive MSCs experienced a decrease in proliferation following IL6 treatment [111], implying that IL6 influence on osteogenesis is complex and dependent on the status of targeted MSCs. In this study, a significant increase in *IL6* expression was observed in LPS-treated gASCs. A similar trend was noted in ST-treated gASCs and hASCs, although not statistically significant. This suggests that ASC response to ST infection influences the secretion of small molecules capable of cross-talk between inflammatory and differentiation pathways.

The transcriptome of hASCs post ST association alludes to a physiological shift in favor of cell survival and proliferation. Under oxidative stress, MSCs display a reduced ability to repair tissue and an increased propensity towards senescence [112, 113]. These conditions decrease MSC capacity for osteogenesis in favor of adipogenic commitment [113]. Through upregulation of redox mediators, hASCs appear to respond and mitigate oxidative stress, helping to insure cell viability, multipotency, and promote immune suppression. As versatile immune privileged cells, MSCs presented with microbial challenge may function as a safeguard, contributing to an anti-inflammatory environment, which allows time and an atmosphere conducive for infection clearance by resident phagocytes. However, it is unlikely that hASC response to ST is without physiological consequences, as influence of inflammatory mediators on lineage commitment appears to prime hASCs towards a pro-osteogenic phenotype.

The direct association of ASCs with ST influences key modulators of trilineage differentiation. We illustrate that, as the pathways dictating MSC response to injury, microbial products and inflammation overlap with the regulation of cellular differentiation, exposure to bacteria alters lineage commitment. However, the extent and mechanisms responsible for eliciting changes in MSC differentiation due to microbial interaction have yet to be fully described. As ASCs are currently in contention as a powerful model for understanding the complex interaction of pathogenesis, inflammation and calcification, further investigation into the complex relationship between bacterial association and ASC physiology is imperative, as we have established that this association results in distinctive changes to MSC physiology.

## Acknowledgments

We thank Majlis Amanah Rakyat (Malaysia) and the Jastro Shields Research Fellowship, and the USDA-CREES W2171 Regional Research Project for funding a portion of the work described in this paper (NMF).

