## Supplementary material for "*Salmonella* Enhances Osteogenic Differentiation in Adipose-Derived Mesenchymal Stem Cells": supp table 1

### qRT-PCR primer sequences for MSC differentiation markers

| Purpose | Gene | Species | | Sequence 5'to 3' | Product size (bp) | Anneal T (°C) | Source |
|---|---|---|---|---|---|---|---|
| ICG | GAPDH | Human | F | ACCACAGTCCATGCCATCAC | 452 | 60 | IDT ReadyMade Primers |
| | | | R | TCCACCACCCTGTTGCTGTA | | | |
| | | Goat | F | TTGTGATGGGCGTGAACC | 127 | 60 | [84] |
| | | | R | CCCTCCACGATGCCAAA | | | |
| Adipogenesis | PPARG | Human | F | TCCATGCTGTTATGGGTGAA | 193 | 60 | [85] |
| | | | R | TCAAAGGAGTGGGAGTGGTC | | | |
| | | Goat | F | AAAGCGTCAGGGTTCCACTA | 201 | 60 | [86] |
| | | | R | CCCGAACCTGATGGCGTTAT | | | |
| | FABP4 | Human | F | GCCAGGAATTTGACGAAGTCAC | 88 | 60 | [87] |
| | | | R | TTCTGCACATGTACCAGGACAC | | | |
| | | Goat | F | TGAGATGTCCTTCAAATTGGG | 101 | 60 | [88] |
| | | | R | CTTGTACCAGAGCACCTTCATC | | | |
| Osteogenesis | COL1A1 | Human | F | GCCGTGACCTCAAGATGTG | 183 | 60 | NM_000088.3 |
| | | | R | CCTTGGGGTTCTTGCTGAT | | | |
| | | Goat | F | TGACTGGAAGAGCGGAGAAT | 158 | 60 | NM_001034039.2 |
| | | | R | CCTTGGGGTTCTTGCTGATA | | | |
| | ALPL | Human | F | GGAACTCCTGACCCTTGACC | 85 | 60 | [89] |
| | | | R | TCCTGTTCAGCTCGTACTGC | | | |
| | | Goat | F | CTGACCACTCCCACGTCTTT | 102 | 60 | M18443.1 |
| | | | R | TGAACGGCTTCTTGTCTGTG | | | |
| | SPP1 | Human | F | GCCGAGGTGATAGTGTGGTT | 125 | 60 | J04765.1 |
| | | | R | ATTCAACTCCTCGCTTTCCA | | | |
| | | Goat | F | CCAACAATCGCAGTTTTCACTC | 81 | 60 | [84] |
| | | | R | CAGTCCATAAGCCACACTATCACC | | | |
| Chondrogenesis | SOX9 | Human | F | GGAGCTCGAAACTGACTGGAA | 151 | 60 | [90] |
| | | | R | GAGGCGAATTGGAGAGGAGGA | | | |
| | | Goat | F | ACGCCGAGCTCAGCAAGA | 71 | 60 | [91] |
| | | | R | CACGAACGGCCGCTTCT | | | |
