## Supplementary material for "*Salmonella* Enhances Osteogenic Differentiation in Adipose-Derived Mesenchymal Stem Cells": supp table 2

### qRT-PCR primer sequences for MSC Inflammatory Markers

| Gene | Species | Sequence 5'to 3' | | Product size (bp) | Anneal T (°C) | Source |
|---|---|---|---|---|---|---|
| IL6 | Human | F | GGTACATCCTCGACGGCATCT | 81 | 60 | [92] |
| | | R | GTGCCTCTTTGCTGCTTTCAC | | | |
| | Goat | F | AAGAATGACACCACCCCAAG | 195 | 60 | NM_001285640.1 |
| | | R | GCATCCATCTTTTTCCTCCA | | | |
| PTGS2 | Human | F | TTCAAATGAGATTGTGGGAAAATTGCT | 305 | 60 | [93] |
| | | R | AGATCATCTCTGCCTGAGTATCTT | | | |
| | Goat | F | AGGCTTCACTTGACCAGAGC | 182 | 60 | JN793538.1 |
| | | R | TACCAGAAGGGCGGGATACA | | | |
| NFKB1 | Human | F | ATGGCTTCTATGAGGCTGAG | 128 | 60 | [94] |
| | | R | GTTGTTGTTGGTCTGGATGC | | | |
| | Goat | F | ACCCTATGAGCCAGAGTTT | 171 | 60 | [95] |
| | | R | AAGGCATTGTTCAGTATCC | | | |
| TGFB1 | Human | F | TACTACGCCAAGGAGGTCAC | 242 | 60 | [96] |
| | | R | GCTGAGGTATCGCCAGGAAT | | | |
| | Goat | F | CACCCGCGTGCTAATGGT | 101 | 60 | [97] |
| | | R | CTCGGAGCTCTGATGTGTTGAA | | | |
| IL8 | Human | F | ATAAAGACATACTCCAAACCTTTCCAC | 102 | 60 | [98] |
| | | R | AAGCTTTACAATAATTTCTGTGTTGGC | | | |
| | Goat | F | TTCCAAGCTGGCTGTTGCTCTCTT | 103 | 60 | [99] |
| | | R | GCATTGGCATCGAAGTTCTGTACTC | | | |
