## Supplementary figures and images for "*Salmonella* Enhances Osteogenic Differentiation in Adipose-Derived Mesenchymal Stem Cells"

### supp fig 1

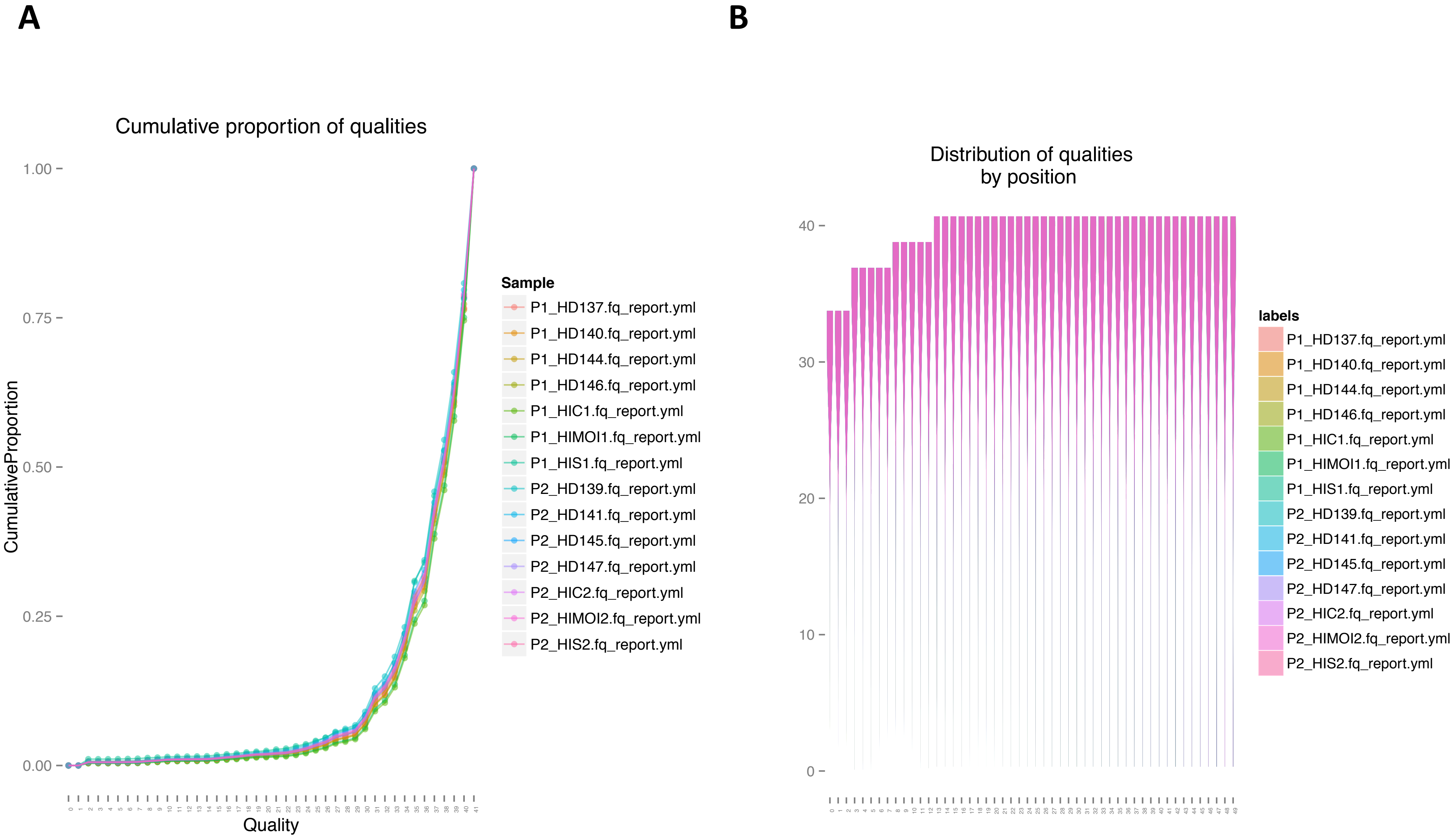

### supp fig 2

A. Caco-2

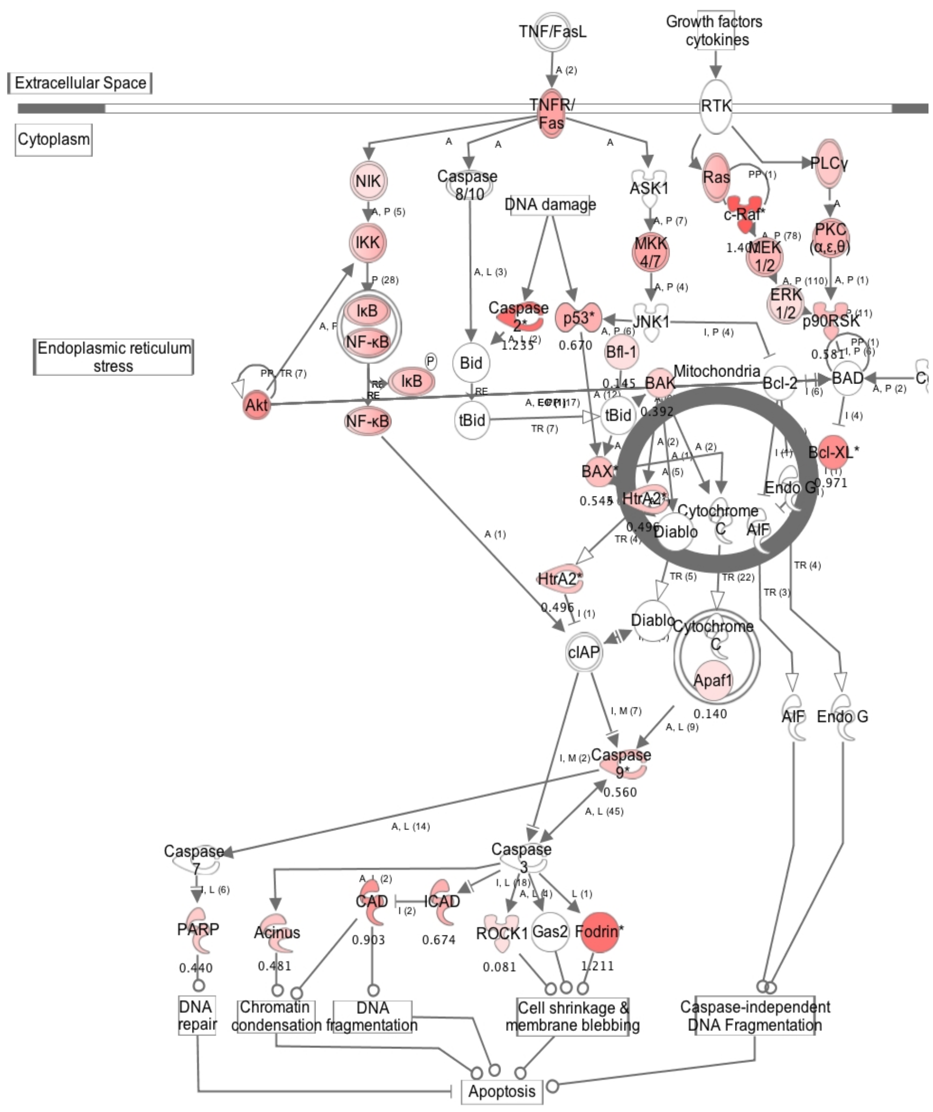


B. hASC

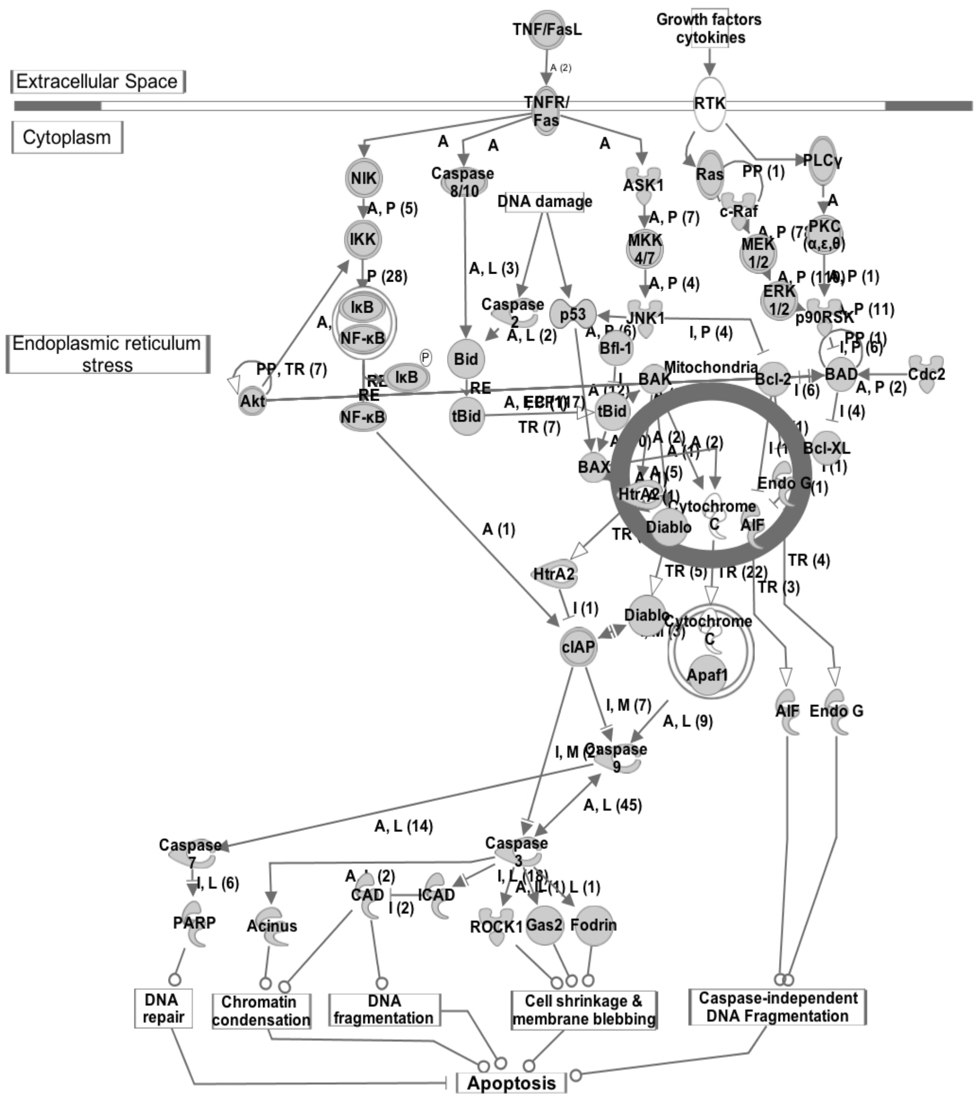

### supp fig 3

#### A. Caco-2

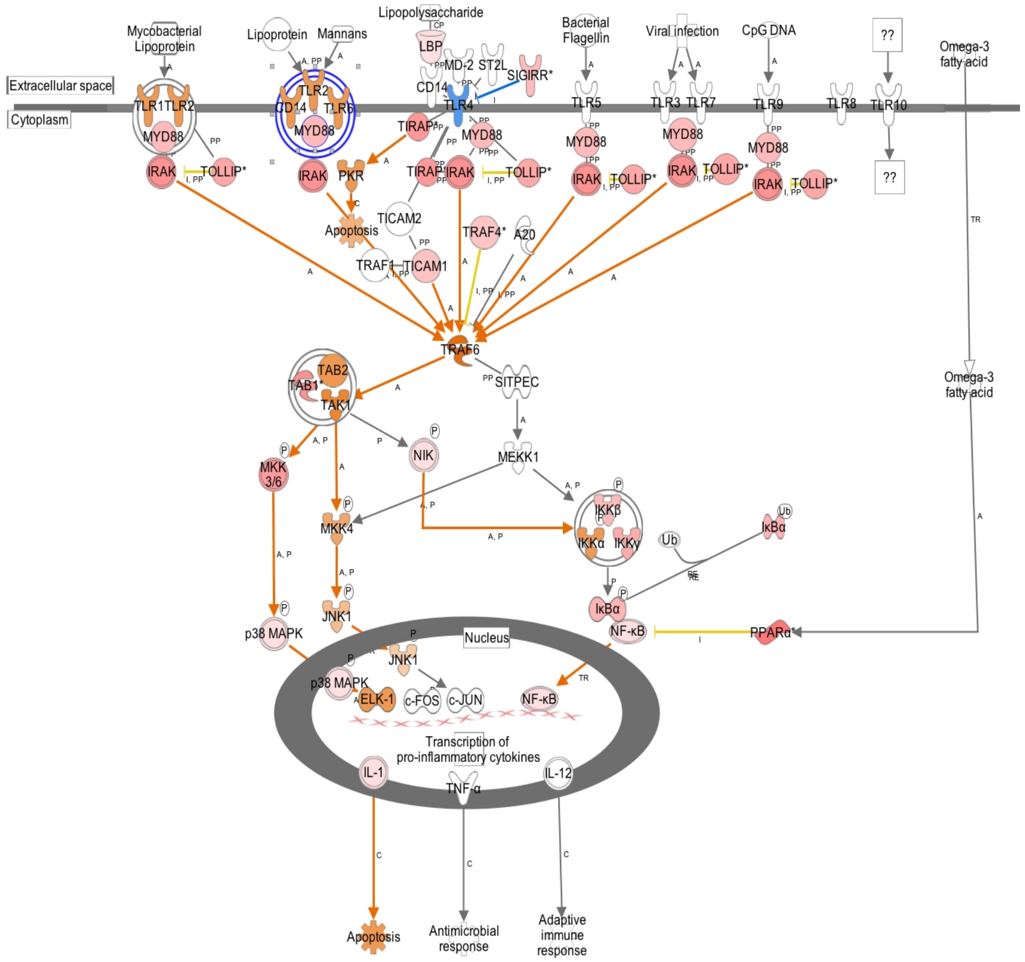


#### B. hASC

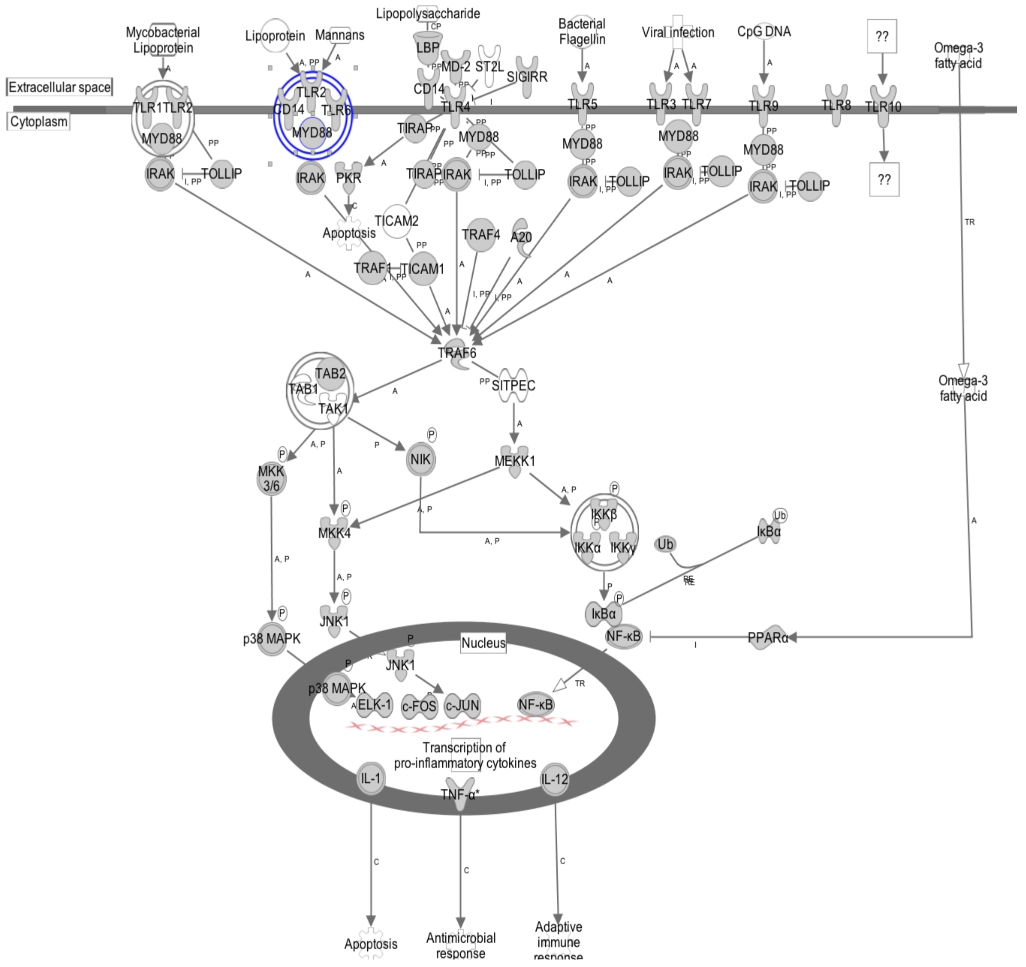

### supp Fig 4B

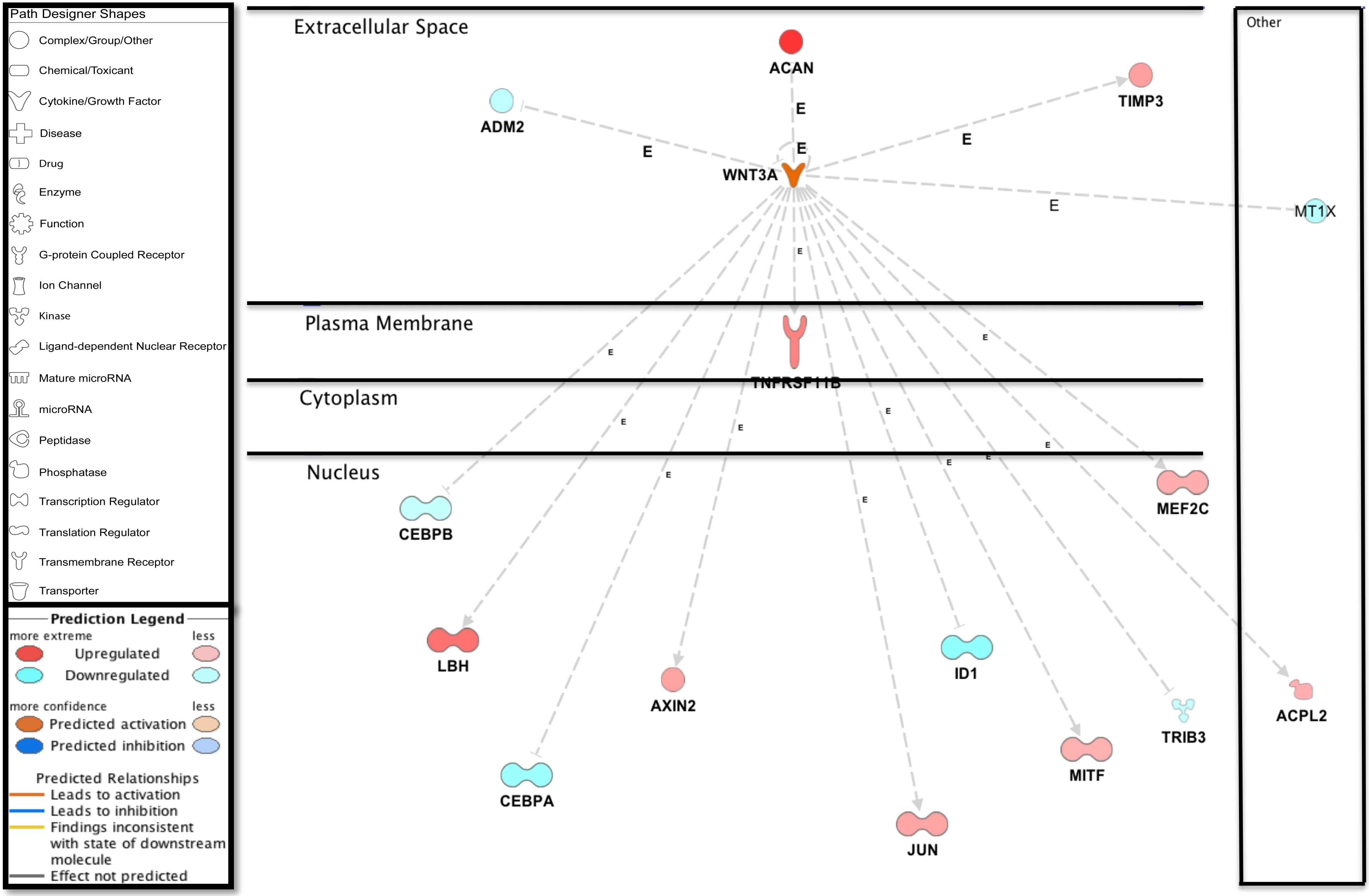

### supp Fig 5

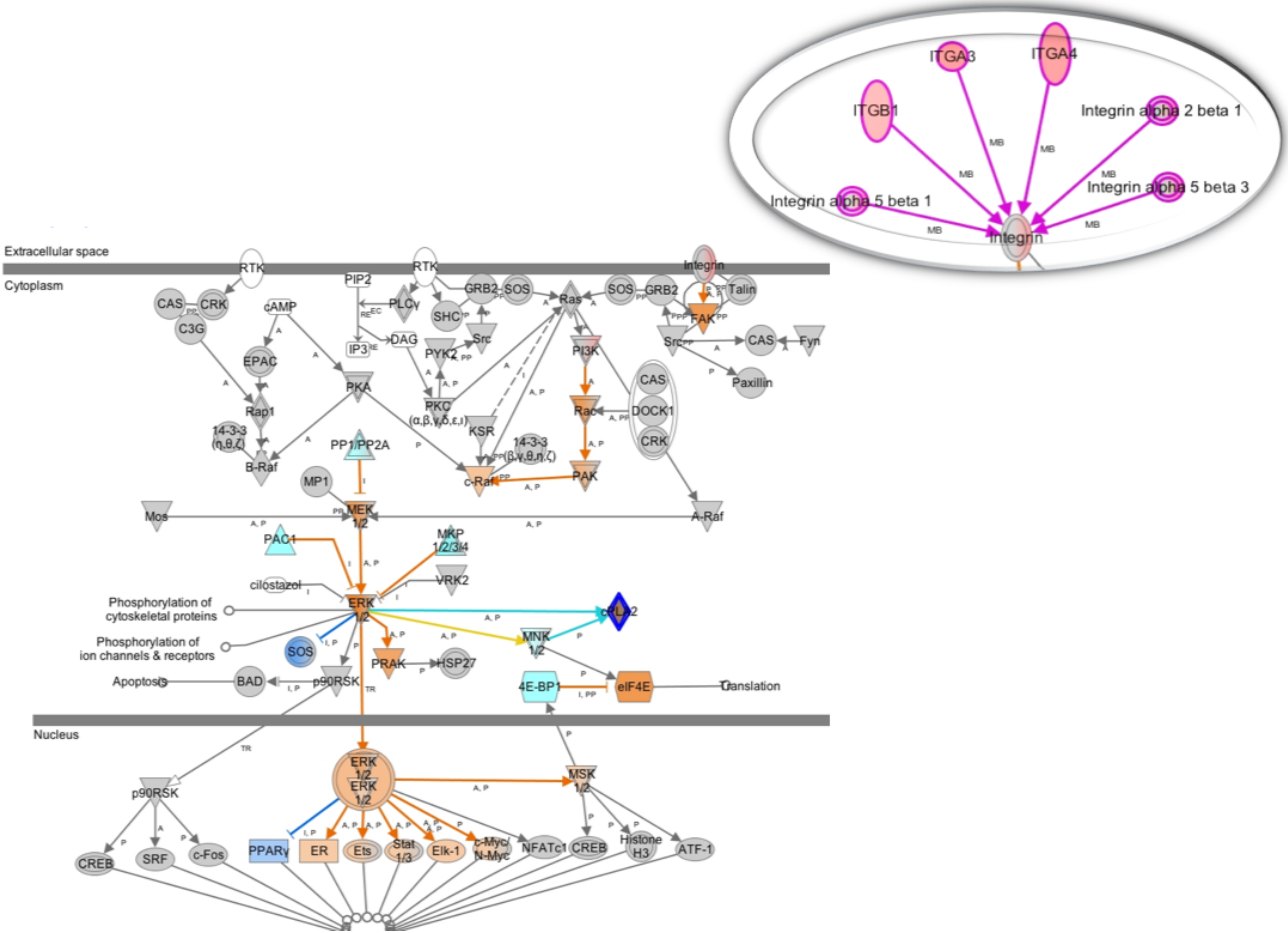

### supp Fig 6

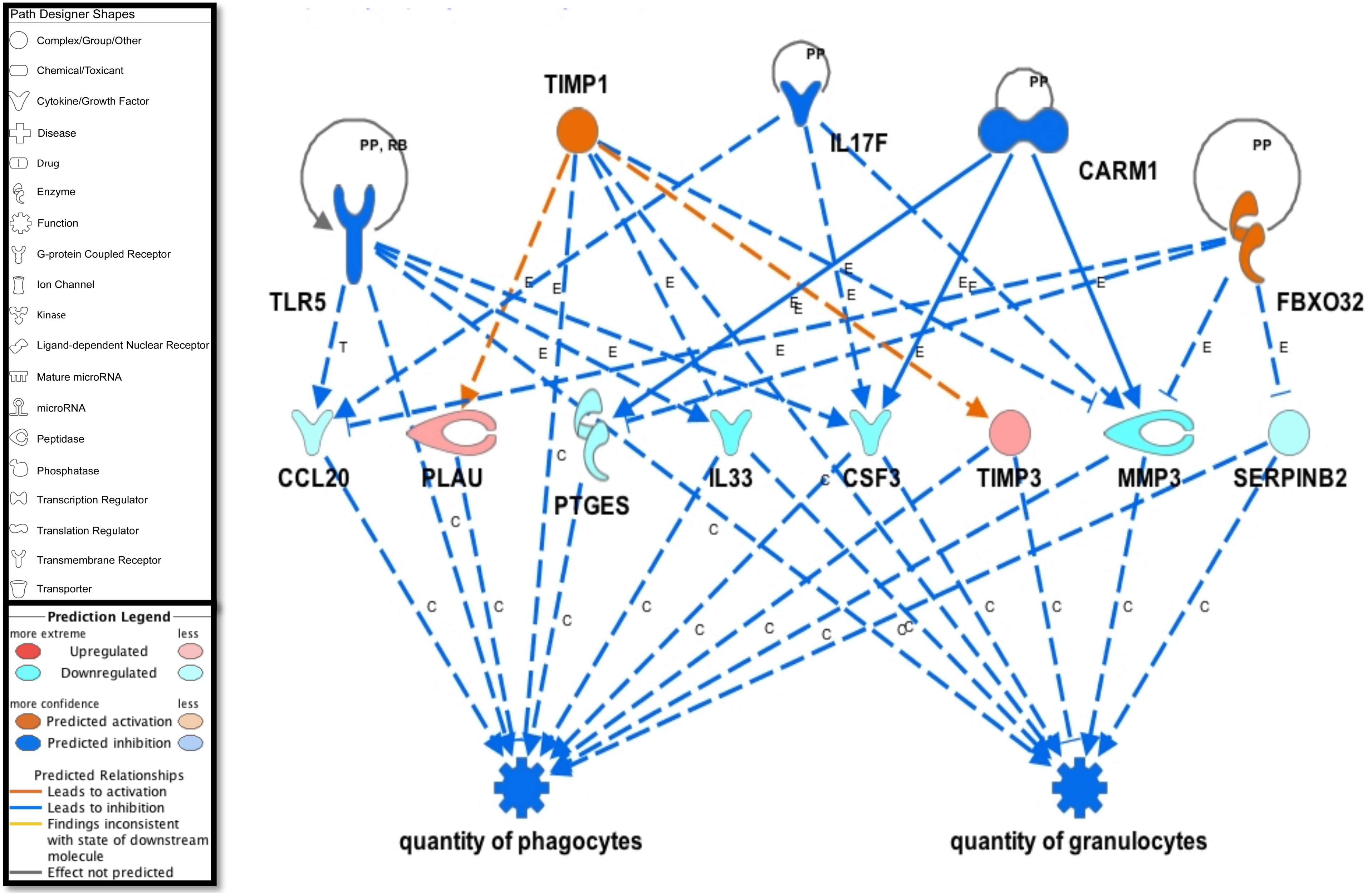

### supp Fig. 4A

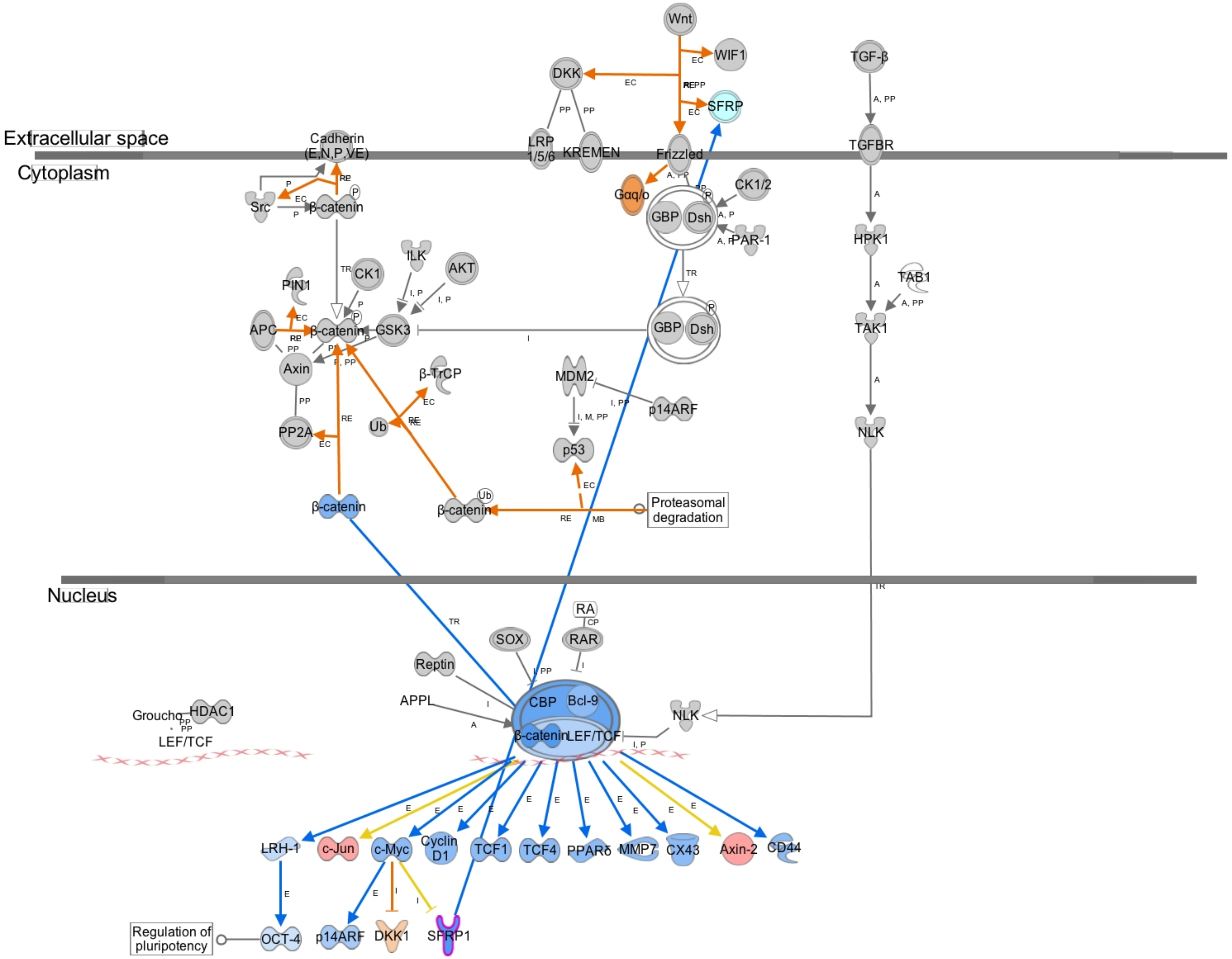
